# Response to divergent selection on meiotic recombination in Saccharomyces cerevisiae

**DOI:** 10.64898/2026.04.02.716186

**Authors:** Xavier Raffoux, Xanita Saayman, Walaa A. Abuelgassim, Thomas Maret, Anthony Venon, Fabrice Dumas, Lorenzo Tattini, Olivier C. Martin, Gianni Liti, Matthieu Falque

## Abstract

Meiotic recombination is a key driver of evolution in sexually reproducing species, reshaping genetic diversity by generating novel allelic combinations. The rate of recombination varies substantially across living organisms depending on *cis*- or *trans*-acting genetic elements, as seen in many species, including the yeast *Saccharomyces cerevisiae*. Here, we report on an experimental evolution-based study to better understand the factors shaping this natural variation. Starting with a genetically diverse population of *S. cerevisiae*, we have carried out recurrent divergent selection on recombination rate using a fluorescence-based sorting approach in four independent lineages. After ten generations, we observed an average response of recombination rate of +28% after positive selection and −24% after negative selection, within the interval used for selection. In the adjacent region, however, we observed a weaker response in the opposite direction, and no response in four other unlinked genomic regions. Whole-genome sequencing of individuals selected for high recombination revealed mixed outcomes in the four independently evolved lineages for high genome-wide recombination rates. However, all four lineages showed selection for high recombination locally, with particular haplotypes heavily favored and sequence- or structural variation-based heterozygosity selected against within the selection interval. Overall, this experimental evolution approach provides original and useful insights into the evolvability of the meiotic recombination rate and the associated genetic determinants.

## Introduction

Meiosis is considered one of the major evolutionary innovations of eukaryotes. The associated genetic recombination is at the heart of many evolutionary processes, both under natural selection (Burt and Bell 1987) and in the genetic improvement of domesticated species (Hill and Robertson 1966; Tourrette *et al*. 2019). Crossovers (COs) occurring during prophase of the first meiotic division have two major consequences: they generate a physical link between homologous chromosomes, which allows a balanced segregation of homologs at anaphase I, required for the viability of the gametes, and they shuffle the parental haplotypes to produce novel allelic combinations in the recombinant gametes. During meiotic prophase I, a large number of programmed DNA double-strand breaks are produced on the chromatids. They are then repaired by several possible pathways which use the homolog as a template — leading to the formation of a minority of COs, which are reciprocal exchanges of genetic material between homologs, and a majority of non-crossovers, which are non-reciprocal short tract exchanges producing gene conversions (GCs), or which use the sister chromatid as a template with no genetic consequence. Two distinct pathways have been demonstrated to exist in many organisms for CO formation (Higgins *et al*. 2004; Housworth and Stahl 2003; Falque *et al*. 2009), one of them being sensitive to the phenomenon of interference that reduces the probability of two COs occurring close to each other (Sturtevant 1915), and the other one being insensitive to interference.

CO number and distribution along chromosomes are strongly regulated (Pan *et al*. 2011). The number of COs per chromosome usually ranges between one and three, and is similar across species, in spite of very large differences in physical chromosome sizes (Fernandes *et al*. 2017). The distribution of COs along chromosomes, often referred to as the CO landscape, is usually uneven, e.g. in yeast (Mancera *et al*. 2008), which is due to a combination of different factors, including interference (Morgan *et al*. 2024). Large pericentromeric regions are almost void of COs in many organisms (Haenel *et al*. 2018), and some very small regions can have high densities of COs (hotspots). CO regulation shows several similar features in various species, see review in (Dluzewska *et al*. 2018). Among them, (1) CO assurance (Jones 1984) ensures that at least one CO occurs per bivalent during meiosis, allowing faithful chromosome segregation, (2) CO homeostasis (Martini *et al*. 2006) keeps the rate of COs almost constant even if the number of double-strand breaks is reduced, (3) CO interference (Sturtevant 1915; Morgan *et al*. 2024) leads to COs being more regularly spaced than if they were placed at random, (4) heterochiasmy (Sardell and Kirkpatrick 2020) leads to different levels and landscapes of recombination in male vs female meiosis, and (5) natural genetic variation of CO rate distribution is observed within many species (Kong *et al*. 2010; Bauer *et al*. 2013; Brekke *et al*. 2023; Johnston 2024). The genetic determinants, molecular mechanisms, as well as mechanistic or evolutionary drivers or trade-offs underlying such diversity of CO-related phenotypes are still partially unknown, although many studies have addressed these questions (see review in Johnston, 2024). Finally, the recombination rate (hereafter noted RR) also shows some phenotypic plasticity, for instance as a response to variation in temperature in Arabidopsis (Lloyd *et al*. 2018) and in yeast (McNeil *et al*., 2025) or nutritional state in Drosophila (Novak *et al*. 2026) and in yeast (Abdullah and Borts 2001).

There are two types of genetic determinants of CO formation: *cis* DNA sequences that modulate RR locally in their vicinity and thus change the recombination landscape, and some *trans* factors that affect RR genome-wide. Both contribute to the natural variability of RR, and we estimated their respective contributions to 38% and 17% of the variance of RR in *S. cerevisiae* (Raffoux *et al*. 2018a). *Cis* effects may involve local heterozygosity (Dluzewska *et al*. 2018). For significant sequence divergence, recombination decreases as heterozygosity increases (Raffoux *et al*. 2018a), as expected if strong divergence suppresses COs through the mismatch-repair system (Hunter *et al*. 1996). In addition, Raffoux *et al*. (2018a) estimated RRs for the four strains that are the founder parents of the SGRP-4X population used in the present study, and observed up to 2.8-fold differences in RR among them, suggesting the presence of alleles of opposing effects at recombination modifier loci. Recombination QTLs acting in *trans* have also been detected in several species (Pan *et al*. 2017; Petit *et al*. 2017; Ziolkowski *et al*. 2017). Moreover, RR could be increased in Arabidopsis and in a few crop species by knocking out anti-CO genes (e.g. FANCM, FIGL1 or RECQ4), overexpressing pro-CO genes (e.g. HEI10), or changing the ploidy context in Brassicaceae, see review in Blary and Jenczewski (2019).

Although CO formation is highly regulated, the evolutionary pressures behind such regulations remain unclear. There must be at least one CO per bivalent for gametes to be viable, and there seems also to be a soft upper limit to CO number in most species studied (Fernandes *et al*. 2017), but the underlying selective pressures remain hypothetical. Recombination is strongly associated with the occurrence of sex, so the fitness cost of recombination itself (crossovers) may not always be easily distinguished from that of sex (segregation) in experimental evolution approaches, as reviewed in Parée and Teotónio (2025). Sex involves spending both time and energy, and has a ‘twofold cost’ because the females transmit their genes at only half the rate of asexual equivalents (Bell 1982). Recombination may also increase or decrease fitness (Charlesworth and Barton 1996): it can break beneficial allelic combinations built up by selection, producing the ‘recombination load’ (Otto and Lenormand 2002), e.g. in *D. melanogaster* where decreased recombination led to higher female fecundity (Barton 2010). Too many COs may also be mutagenic (Liu *et al*. 2017) or could be mechanically deleterious for meiosis, although evidence for this is still lacking. Yet despite these costs, sex and recombination are maintained in the great majority of species, which is referred to as the ‘paradox of sex’ (Otto and Lenormand 2002). On the other hand, shuffling alleles through recombination can be beneficial for adaptation by increasing the genetic variance (Barton 2010). Experimental evolution of yeast populations revealed that sexual reproduction favored adaptation to a new harsh environment, whereas no effect was observed in a new benign environment (Goddard *et al*. 2005), and that sex both speeds up adaptation and allows natural selection to more efficiently sort beneficial from deleterious mutations (McDonald *et al*. 2016). Altogether, the relationship between recombination and fitness is difficult to study because it is mostly indirect: COs do not directly change fitness (except a small number of COs required to keep euploidy through meiosis), but they modify the potential to exploit standing genetic variation to produce new haplotypic combinations, which may be beneficial (or deleterious) for fitness. Recombination modifier genes are thus expected to be able to evolve as a response to such indirect selection pressures (Dapper and Payseur, 2017). These authors review many theoretical works indicating that indirect selection strongly depends on the status of linkage disequilibrium in the population, producing coupling or repulsion situations, which may be driven by different genetic mechanisms among which epistasis and genetic drift can play important roles. They also note that even though many theoretical studies have investigated possible effects of such indirect mechanisms on the evolution of recombination, there is a lack of cross-feed between these theoretical works and empirical ones describing the observed diversity of RR or its evolvability in natural or laboratory populations. In the particular case of agricultural species, the domestication syndrome can involve many loci, potentially leading to an indirect increase of recombination as confirmed by experimental observations in several livestock species (Bursell *et al*., 2025). Finally, CO interference can also be an important player for the adaptive consequences of recombination, and thus for indirect selection of recombination modifiers, since it lowers the probability that two close COs bring together three favorable alleles linked in repulsion phase (Otto and Payseur 2019). On the contrary, there is little evidence for a direct relationship between RR and fitness, and thus whether recombination rate experiences direct selection in natural populations (Ritz *et al*., 2017). Nonetheless RR shows intraspecific diversity, and many adaptive processes have been found to be associated with the evolution of recombination modifier genes in theoretical (Gandon and Otto 2007) as well as empirical (Korol and Iliadi 1994; Aggarwal *et al*. 2015) studies.

In this context, applying direct recurrent artificial selection on recombination rate should make it easier to investigate the evolvability of recombination, and to shed light on the mechanisms at work in its response to selection, which is our focus in the present work. However, selecting on RR is challenging because of the difficulty to measure recombination phenotypes. COs may be detected by studying recombination between genetic markers in linkage mapping experiments (Bauer *et al*. 2013) or by direct cytological observations. In the latter case, one can measure CO positions by visualizing late recombination nodules using electron microscopy (Anderson *et al*. 2003) or by immunofluorescence using antibodies against different proteins that concentrate as foci at the location of COs (Anderson *et al*. 1999; Chelysheva *et al*. 2012). In all cases, these methods are destructive, and they are difficult to bring to a high throughput as required for experimental evolution.

Nevertheless, direct selection experiments on RR have been reported a long time ago in Drosophila, based on the segregation of morphological genetic markers. After six generations of positive or negative selection on CO rate in the interval between loci ‘dichaète’ and ‘sepia’, no response to selection was first observed (Gowen 1919), but later on, during 30 generations of selection for low CO rate in the interval between ‘white eyes’ and ‘miniature wings’ loci, a continuous reduction of RR was observed (Detlefsen and Roberts 1921). In the same paper, eight generations of selection for high RR in the same interval also produced a reduction of recombination, which was interpreted by the authors as a positive response, selection producing many double COs. Several similar experiments have been performed in other genomic regions of Drosophila (Parsons 1958; Chinnici 1971a; Kidwell 1972; Abdullah and Charlesworth 1974; Charlesworth and Charlesworth 1985a), which confirm that recurrent artificial selection for high or low recombination rate most often results in a genetic response. Further studies on the genetic architecture of the response showed that it is polygenic (Chinnici 1971b; Charlesworth and Charlesworth 1985b) and not due to chromosomal rearrangements. In the insect *Tribolium castaneum*, significant response to selection for high RR has also been reported during 10 generations, after which a plateau was reached, and RR was stable for five more generations (Dewees 1975). In the same paper, when selecting for low RR, there was no significant decrease of RR along generations, but the average RR over the 15 generations was significantly lower than the initial value, suggesting a response limited to the first generations. In *Schistocerca gregaria*, the response to selection was significant for low RR but not for high RR, although there was a trend suggesting a weak response in the second case (Shaw 1972). In silkworm, response to selection was obtained for both low and high RR, as cited by Ebinuma and Yoshitake (1981). Apart from insects, a similar experiment has been achieved in Lima bean (Allard 1963), resulting in significant responses to selection for high RR but no response to selection for low RR. In animals and fungi, we did not find any published results on long-term experimental evolution carried out with artificial selection on RR. Although the organisms studied and the type of experimental evolution conditions were very different across these studies, altogether they suggest that a response to artificial selection for RR might be expected within approximately ten generations, which is what we aimed at in the present work.

Such studies on the evolvability of CO number based on experimental direct recurrent selection on RR can provide useful insights on the evolutionary dynamics of recombination modifier genes that can play a crucial role in adaptation from standing variation, particularly in changing environments (Charlesworth 1993). They can also produce evolved populations with diverged recombination rates, which may then be used for experimental evolution with the aim of studying the impact of recombination rate on the dynamics of adaptation after environmental changes. Moreover, such populations may be used to detect signatures of selection by applying sequencing using the ‘evolve and resequence’ approach (Long *et al*. 2015), which could point to recombination QTLs and help identify genes or other genomic elements involved in the control of the quantitative variation of CO rate.

Here, we carried out experimental evolution on a yeast population with high standing genetic variation by applying ten cycles of sexual reproduction with recurrent divergent selection of spores based on their recombination status in a region between two fluorescent markers. We observed a response to selection in the interval where selection was applied and a corresponding inverse response in a neighboring interval. Selecting for recombination over eight cycles led to an increase in homozygosity within the selection interval on both the structural and sequence levels. Genome-wide CO analysis further revealed a global increase in CO rate in two of four lineages without impacting the number of non-crossovers. These results collectively suggest that we successfully selected for high-recombination on both the *cis* and *trans* levels in yeast. They also provide details on the dynamics and chromosomal location of RR changes which allow us to formulate hypotheses about the drivers and constraints shaping the evolvability of recombination.

## Materials and Methods

### Biological material

The original SGRP-4X population (Cubillos *et al*. 2013) was obtained by inter-crossing for 12 generations four representative *S. cerevisiae* founder strains: YPS128 [North American (NA)], DBVPG6044 [West African (WA)], Y12 [Sake (SA)], and DBVPG6765 [Wine/ European (WE)]. The bi-fluorescent strain Mat-a SK1-VI_C1Y2 ho::hygMX ura3::kanMX used to cross to the SGRP-4X population was constructed as previously described (Raffoux *et al*. 2018b). Details on the genomic positions of the two fluorescent markers used to select on recombination are given in Suppl Text section 1. For recombination phenotyping, we also used the three tri-fluorescent tester strains SK1-I_R2C3Y4, SK1-VI_C1Y2R3, and SK1-VI_R3Y4C5. Details of constructs, genomic positions, and physical distances between markers for all fluorescent markers used in this work are given in Suppl Table S1.

### Culture Media and Drugs

Solid SPOR medium was prepared with 0.25 % yeast extract, 0.1 % glucose, 1 % potassium acetate, and 20 g/L BD Bacto Agar (BD-214030). Nourseothricin was used at a final concentration of 30 mg/L, G418 at 100 mg/ L, chloramphenicol at 50 µg/mL, and hygromycin B at 200 µg/mL.

### Crossing the SGRP population with SK1-VI_C1Y2

To produce the initial population (G0) of our evolution experiment, we crossed spores from the SGRP-4X population with haploid vegetative cells of the SK1-VI_C1Y2 bi-fluorescent strain. First, the SGRP-4X population was sporulated and spores isolated as described in Suppl Text section 2. Then, the spores were germinated and mixed with vegetative cells of the haploid Mat-a bi-fluorescent strain SK1-VI_C1Y2, and diploids originating from crosses between a SGRP-4X spore (Ura+) and a tester cell (natMX) were selected on specific medium (see Suppl Figure S1) as detailed in Suppl Text section 3. Selected diploids were stored and used as the generation zero (G0) to start the selection experiments.

### Selection of spores depending on their fluorescence status

Isolated spores from the G0 diploid population were obtained with the same protocol as presented above and in Suppl Text section 2. The spores suspension was then analyzed with a MoFlo ASTRIOS Fluorescence-Activated Cell Sorter (FACS; Beckman-Coulter, Villepinte, France) and the associated software Summit for manual investigations, or with the CAYSS R package (Raffoux and Falque 2024) for routine analyses. The FACS was first used to select spores from vegetative cells based on their size and granularity, and then to sort spores having the desired fluorescent marker combination as detailed in Suppl Text section 4. Fluorescence intensity classes for spore selection were chosen depending on the direction of the selective pressure to be applied: selection of spores recombinant between the fluorescent markers (Sel+; Figure 1A), selection of spores non-recombinant between the markers (Sel-; Figure 1B), or selection of all events identified as spores, regardless of their recombination status between the markers (Sel=; Figure 1C).

**Figure 1:**
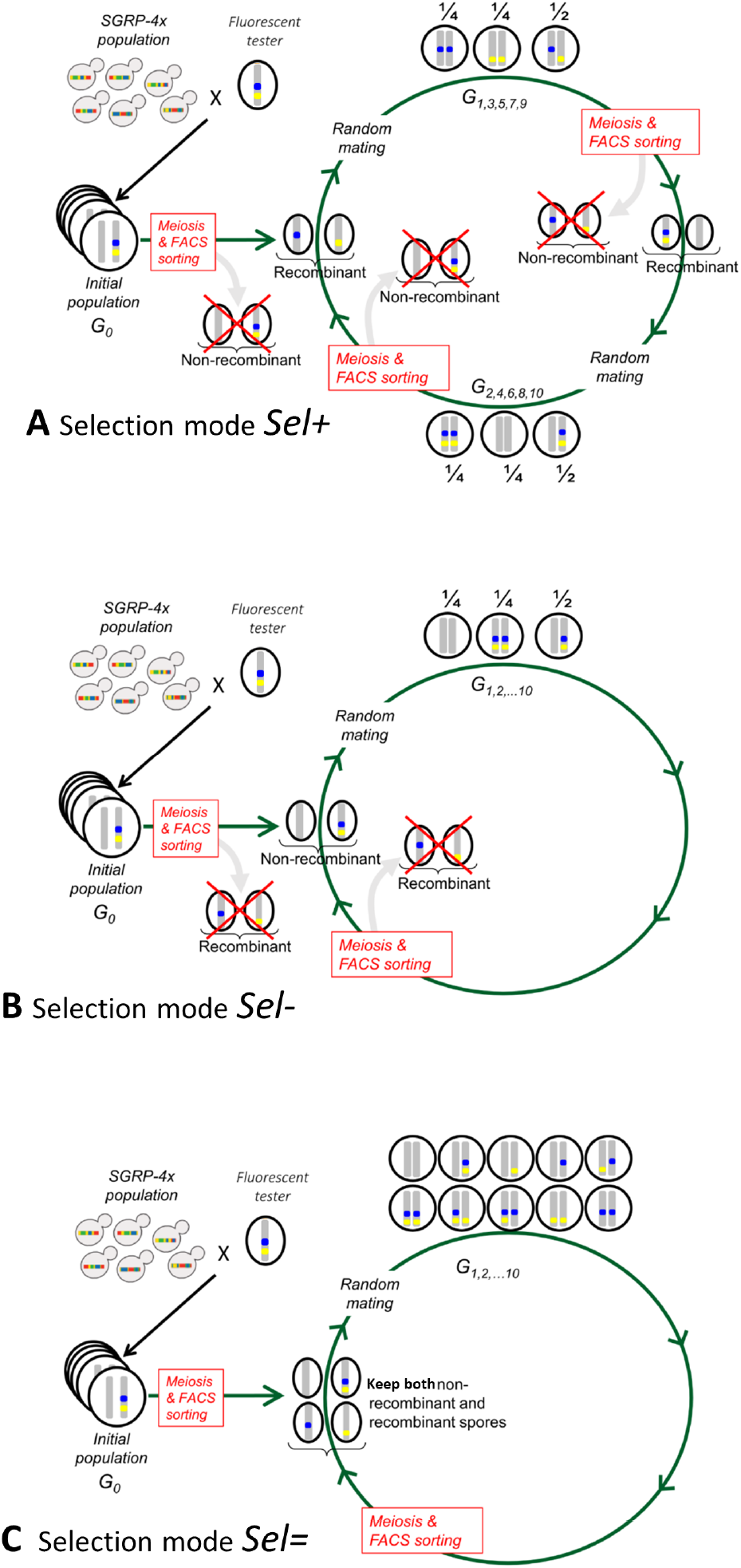
Details of the three modes of selection applied to spores. In Sel+ experiments, only recombinant spores are selected by FACS-sorting (A), in Sel-, only non-recombinant spores are selected by FACS-sorting (B), and in Sel= controls, all spores are kept (C), and in all three selection regimes, a total of 300000 spores are sorted for the next generation. Note that in Sel+, two selection cycles are represented on the sketch: odd generations on the top side of the circle, and even generations on the bottom side of the circle. In contrast, for Sel- and Sel=, one single cycle of selection is represented in the sketch.

In the following, we will represent the phenotype of the two fluorescent markers C and Y by “[MN]” where M is “C” if the marker yECerulean is present or “+” if it is not present, and N is “Y” if the marker Venus is present, and “+” if it is not present. So in even generations of Sel+ experiments, 150,000 [C+] recombinant spores and 150,000 [+Y] recombinant spores were selected (left side of Figure 1A), in odd generations of Sel+ experiments, 150,000 [++] recombinant spores and 150,000 [CY] recombinant spores were selected (right side of Figure 1A), in Sel-experiments, 150,000 [CY] non-recombinant spores and 150,000 [++] non-recombinant spores were selected (Figure 1B), and in Sel= experiments, we sorted a total of 300,000 cells from the four fluorescence classes [CY], [++], [C+], [+Y] (Figure 1C), ensuring to sort equal numbers of [CY] and [++] spores, and equal numbers of [C+] and [+Y] spores. Using these procedures for each of the three selection modes (Sel+, Sel-, Sel=), we sorted four biological replicates (noted A, B, C, D) from the initial G0 population, leading to 12 independently evolved populations for each generation, hereafter noted Gn Sel+ A, Gn Sel+ B, Gn Sel+ C, Gn Sel+ D, Gn Sel-A, Gn Sel-B, Gn Sel-C, Gn Sel-D, Gn Sel= A, Gn Sel= B, Gn Sel= C, Gn Sel= D, where n is the index of the generation. To produce the G2 populations, G1 populations were then again sporulated, and spores FACS-sorted following a similar procedure as from G0 to G1 (details in Suppl Text section 5). This recurrent selection was conducted during 10 generations (noted G1 to G10). Note that as seen before, in the case of Sel+ experiments, fluorescence classes to be selected for recombinant and non-recombinant spores are not the same for even or odd generations.

### Pool-phenotyping recombination in populations

To phenotype recombination at each generation and for each of the 12 populations (replicates A, B, C, D of selection modes Sel+, Sel-, Sel=), 50,000 non-fluorescent spores were sorted in three different Petri dishes containing 100 µL of vegetative cell suspension (at plateau phase) of each of the three haploid tester strains SK1-I_R2C3Y4, SK1-VI_C1Y2R3, and SK1-VI_R3Y4C5 (Suppl Figure S2).

As the Venus (Y) fluorescent marker can exhibit 3 fluorescence levels as mentioned above, depending on the selection mode, the positions of the gates used to select the non-fluorescent [++] spores to be crossed with the testers were not always the same (see details in Suppl Text section 6). As a consequence, recombination values are comparable only between odd generations or between even generations of Sel+ experiments, but values measured at an odd generation of Sel+ may not be compared to values measured at an even generation. On the other hand, recombination values phenotyped in Sel-, Sel=, and even generations of Sel+ experiments are comparable to each other. Moreover, in odd Sel+ generations (top of Figure 1A), non-fluorescent spores must be produced via recombination happening in the marker interval during sporulation, whereas in even generations of Sel+ (G0 and bottom of Figure 1A) as well as in all generations of Sel-(Figure 1B) and Sel= (Figure 1C), non-fluorescent spores can also be produced without recombination. So in odd Sel+ generations, an extra selection for high recombination is applied during the phenotyping process, whereas this does not happen in other cases.

Diploid cells were then sporulated for 10 days, and spores were isolated (see Suppl Text section 6) and analyzed with a CytoFLEX flow cytometer, the associated software CytExpert (Beckman-Coulter, Villepinte, France) for manual analyses, and the dedicated R package CAYSS (Raffoux and Falque 2024) as described in Suppl Text section 6. RR values are expressed as the fraction of recombinant spores, so without unit. The coefficient of coincidence (CoC) is calculated as the ratio between the observed frequency of double COs (one in each of two adjacent intervals of each tester) and the expected frequency of such double COs in the absence of interference, the latter being estimated as the product of RRs of both intervals (i.e., assuming CO events are independent of each other). So CoC equals one in the absence of interference, and is below one in case of positive CO interference.

### Phenotyping recombination of isolated individuals from G8

The G8 populations were used to FACS-sort bi-fluorescent diploid cells which were then spread on solid medium to isolate individual colonies (see details in Suppl Text section 7). For each of the 12 populations, 288 single cells were sorted. Ninety-six colonies were picked for each of the four biological replicates of the Sel= populations, and 192 colonies for each of the four biological replicates of the Sel− and Sel+ populations.

Out of the 1,920 isolated colonies, some were discarded for different reasons: not enough spores produced, no segregation due to homozygous marker as expected in some cases (see Suppl Figure S3), non-Mendelian single-locus (Suppl Figure S4) or two-locus (Suppl Figure S5 and S6) segregation. Detailed procedures and numbers of discarded strains at each step are given in Suppl Text section 7. FACS-sorted [CY] vegetative diploid cells from G8 Sel= populations can carry two possible marker haplotypes: CY/++ or C+/+Y, each with a probability of 0.5 (Suppl Figure S3). As the cytometry analysis software was parameterized to compute RRs with CY/++ marker configurations, we measured in Sel= strains 44 RR values greater than 0.5 corresponding to C+/+Y configurations and 57 values lower than 0.5, corresponding to CY/++ configurations. This proportion of 0.44 was not significantly different from the expected value of 0.5 (exact binomial test, *p*-value = 0.23). However, we also obtained RR values greater than 0.5 for 28 Sel+ strains and for one Selstrain, which does not match with usual genetic expectations (Suppl Figure S3). Moreover, two Sel+ strains produced RR values very close to 0.5 (0.488 and 0.491). These 31 strains were discarded from the analysis, and the one with RR=0.491 was selected for whole-genome sequencing (see below).

### Tetrad dissection and whole-genome sequencing

Individual strains isolated from Sel+ G8 populations and phenotyped (as described above) were then chosen for tetrad sequencing. Overall, we selected Sel+ individuals with the lowest RR (hereafter denoted [low]) and Sel+ individuals with the highest RR (hereafter denoted [high], see Suppl Table S2) of each biological replicate.

Each individual strain was inoculated overnight in 10 ml YPD at 30 °C, shaking at 220 rpm. The following day, overnight cultures were centrifuged, washed in 10 ml ddH_2_O, and resuspended in 2 % KAc at 1 OD600/ml, and left shaking at 23 °C. After one week, samples that were sporulated were tetrad dissected using standard procedures (see below). Of the 16 individual samples, 2 did not sporulate (one from biological replicates A [high] and one from biological replicates C [low]). For tetrad dissections: 400 µL of the culture was removed and centrifuged (1,000 g, 2 min). Pellets were resuspended in 100 µL dissection buffer (1 M sorbitol, 0.2 mM EDTA, 10 mM sodium phosphate) containing 5 µL zymolyase (UZ1000-A, 5 mg/mL), and incubated for 20 minutes at 37 °C. An additional 400 µL of dissection buffer was added, and samples were placed on ice. After cooling down, 20 µL was used to perform tetrad dissections with the Singer Tetrad Dissection microscope. Plates were grown at 30 °C for 2 days.

A total of 80 segregants from 20 tetrads (9 from [high] strains, 11 from [low] strains) were chosen for whole-genome sequencing. DNA was extracted using the MegaPure kit (MPY80200) using an overnight culture in 5 ml YPD (shaking at 220 rpm, 37 °C) and submitted for whole-genome microbial sequencing with Novogene.

The sequenced tetrads are named according to the sample from which they were taken. For example, “high_A_2.1” refers to the first tetrad dissection of the second sample of the high-recombination lineages ([high]) sorted from Sel+, of biological replicate A. Tetrads that independently went through meiosis but started from the same isogenic sample can be identified by the final number, e.g., high_A_2.1 and high_A_2.2.

### Recombination analysis from tetrad whole-genome sequencing

To reconstruct segregant genotype compositions, diagnostic polymorphisms were identified by first running whole-genome alignment of subtelomere-masked genomes of all four parental genomes, as well as SK1, using progressiveMauve (released 2015-02-13, default parameters) and then extracting polymorphisms unique to each parent. At this point, WA and SK1 unique polymorphisms were merged into one, as they are genetically highly similar. Short read sequences of each segregant were then mapped to the SGD reference genome (S288C, SGD-R64-1-1) using bwa mem (v.0.7.17-r1198-dirty, default parameters) and variants (skipping indels) were called using bcftools (v.1.20). Variants were then cross-referenced with diagnostic parental polymorphisms to identify regions of the genome that could only belong to a single parental genotype (custom scripts, python v.3.12.4). Contiguous tracts belonging to a single parental genotype were then merged, and transition points between genotypes were further refined. To refine transition points: once two adjacent genotypes were known, a second round of identifying diagnostic polymorphisms between the two adjacent genotypes was performed, followed by crossreferencing with segregant variants. Transition points were then called as the base pair directly following the last known unique polymorphism.

To identify structural homology between SK1 and the four parental genomes within the chromosome VI selection interval, mVista (LAGAN, RankVISTA probability threshold p = 0.5) was used to determine conserved regions (CNS) between each pair of genomes. We further filtered conserved regions to require a minimum conservation of 85 % and a minimum region size of 1 kbp, and these were visualized using Python.

CO counts were performed manually using genotype patterns, and GC counts were calculated as the number of contiguous blocks that had at least 2 genotypes present but were not 2:2 between chromatids, as quantified using bedtools (v.2.31.1). To adjust for the presence of regions that are genetically identical between all chromatids (which prevents the detection of COs and GCs), the number, size and location of these blocks were identified for each tetrad using bedtools (v.2.31.1). CO counts were adjusted for any genetically identical blocks on chromosome termini, since CO events within internal genetically identical blocks can be partially inferred by surrounding genotypes. GC counts, on the other hand, were adjusted for all genetically identical blocks.

### Whole-genome sequencing of an isolated diploid strain with an outlier RR value

We used Oxford Nanopore Technologies to whole-genome sequence one diploid strain (1A-E8) isolated from the G8 Sel+ population, which had a RR of 0.491 classifying it as an outlier. Details on DNA extraction, sequencing, and data analysis are given in Suppl Text section 8.

### Checking for possible effects of fluorescent markers on fitness

To assess the extent to which the presence and/or expression of fluorescent marker genes had an effect on cell fitness, G0 cells were sporulated and spores were isolated from vegetative cells as described above. Then, a total of 2,000,000 spores were sorted by FACS (see below) based on their fluorescence status: non-fluorescent [++], yECerulean only [C+], Venus only [+Y], or with both markers [CY]. Sorted spores from each category were plated on solid YPD supplemented with 50 mg/L chloramphenicol and incubated at 30°C for 48 h to allow germination and random mating. Diploid cells were then selected in SD medium lacking uracil and lysine. After overnight growth (10 mL SD–ura–lys in 50 mL tubes, 30°C, 220 rpm), cultures were centrifuged for 5 min at 3 000 rpm, the supernatant was removed, and cells were resuspended in 1 mL ddH_2_O. Fitness measurements were then carried out on these diploids by measuring growth rate, sporulation efficiency, and spore viability as explained as follows:

#### Vegetative growth kinetics

were monitored in 200 µL of cell suspension diluted to an OD_600_ of 0.01 and distributed across the wells of a 96-well microplate to account for possible temperature variation within the plate. The remaining wells were supplemented with sterile YPD as negative controls to control potential contamination. In total, five microplates were prepared. Additionally, 300 µL of each suspension was spread onto four SPOR plates, incubated at 30°C for ten days for subsequent sporulation analysis. Growth kinetics were studied by measuring turbidity every 10 min for 48 h using five custom MiniRead 96-well plate readers (Falque *et al*. 2024). Growth rates were then computed from the growth curves as the maximum growth rate, using the MiniRead R package and calibration procedures associated with the devices.

#### Sporulation rates

were measured after 10 days of sporulation as before, by quantifying the proportions of tetrads and vegetative cells under the microscope, after scraping cells from the solid medium and resuspending them in 10 µL ddH_2_O on a microscope slide.

#### Spore viability

was assessed by sorting single spores by FACS after sporulation of diploids and spore isolation (see section ‘Selection of spores depending on their fluorescence status’). A total of 768 spores from sporulated G0 diploid cells were sorted and deposited on solid YPD in eight one-well rectangular plates (96 spores per plate) for each spore category. Plates were then incubated at 30 °C for 2 days and colonies were counted.

### Fitness assessments of evolved populations at generation eight (G8)

To assess fitness in evolved populations, growth kinetics, sporulation rate, and spore viability were measured as explained before for G0, but from diploid G8 cells coming from each of the three selection modes (Sel+, Sel-, Sel=). For spore viability assessment, a total of 2304 [++] non-fluorescent spores (to limit confounding effects) were sorted from spore preparations of sporulated G8 diploid cells for each of the three selection modes (Sel+, Sel-, Sel=) and deposited onto 24 rectangular YPD plates (96 spores per plate). FACS sorting parameters were set to select spores and discard vegetative cells and debris based on FSC/SSC. Sorted spores from each category (Sel+, Sel-, Sel=) were plated on solid YPD supplemented with 50 mg/L chloramphenicol and incubated at 30°C for 48 h to allow germination and random mating. Fitness measurements were then carried out on these diploids by measuring growth rate, sporulation efficiency, and spore viability as described before.

### Statistical analyses

All data analyses and statistical tests were carried out using R v4.5.0 (R Core Team, 2026) or python v.3.12.4 with several libraries detailed in Suppl Text section 9. In particular, for pairwise comparisons of means, independent *t*-tests were calculated using the stats package from SciPy (Virtanen *et al*., 2020). Multiple comparisons of means were performed by ANOVA using the lm() function of R after checking that variances were homogeneous across treatments and residuals were close to normality, otherwise specific non-parametric tests were used as specified in the text. Contrasts were addressed using post-hoc Tukey tests. In the case of proportions, we used the glm() function with parameter family=binomial(). All multiple tests were corrected using Bonferroni procedure. To quantitatively characterize the distributions of RR among individuals isolated from the evolved populations, symmetry of distributions was tested with the function symmetry.test() from the R package “lawstat” with option=“MGG” (Miao *et al*. 2006), side=“both”, boot=TRUE, B=1000, and q=8/9. The degree of skewness of distributions was quantified using Pearson’s third-order moment coefficient of skewness γ1 (Joanes and Gill 1998).

## Results

### 1. Recombination rate responds to divergent selection

In our divergent selection experiment, we started with an advanced intercross yeast population known as SGRP-4X. SGRP-4X was originally created by recurrently mating four genetically divergent wild founders from the Sake (SA), North America (NA), West African (WA), and Wine/European (WE) clades of *S. cerevisiae*. This highly admixed population results in a representative pool of genetic diversity, which can be used as a powerful tool for mapping quantitative traits (Cubillos *et al*. 2013).

To artificially select for low- or high-recombination, we bulk-mated SGRP-4X with a tester strain in the SK1 background, which contains two fluorescent markers (VI_C1 and VI_Y2) positioned on chromosome VI (Suppl Figure S2). We then recurrently selected for (Sel+) or against (Sel-) spores recombinant within that selection interval VI_C1Y2, based on the segregation of fluorescent signals (Figure 1). At each generation of selection, the recombination rate (RR) was phenotyped by sorting non-fluorescent spores obtained from the diploid population, mating back to fluorescent tester haploid strains, and quantifying RR, expressed as the fraction of recombinant spores, within each interval (Suppl Figure S2) on a population level from the segregation of fluorescent signals in spores. It should be noted that because of this selection for non-fluorescent haploids, only RR values measured in even generations are comparable between Sel+, Sel-, and Sel= experiments (see Figure 1, fully explained in Materials and Methods).

We performed ten sexual generations of selection in four independent lineages (A, B, C, D) using VI_C1Y2 (size: 69.9 kbp) as the selection interval. When phenotyping RR within this same interval, we observed a clear response to directional selection (Figure 2A). The four populations G0, G10 Sel+, G10 Sel=, and G10 Sel-had a significant effect on RR (*p*-value=2.5⨯10^−4^) with all pairwise comparisons significant (*p*-values<0.016) except G0 vs G10 Sel= (*p*-value=0.998). When looking at the evolution of RR across generations (Figure 2A), the curves and their confidence intervals suggest that RR diverged between G0 and G5 or G6. From G6 or G7 on, RR values seem to reach a plateau during which the values of Sel+ and Sel-relative to Sel= were on average +27.6% and −24.5%, respectively.

**Figure 2:**
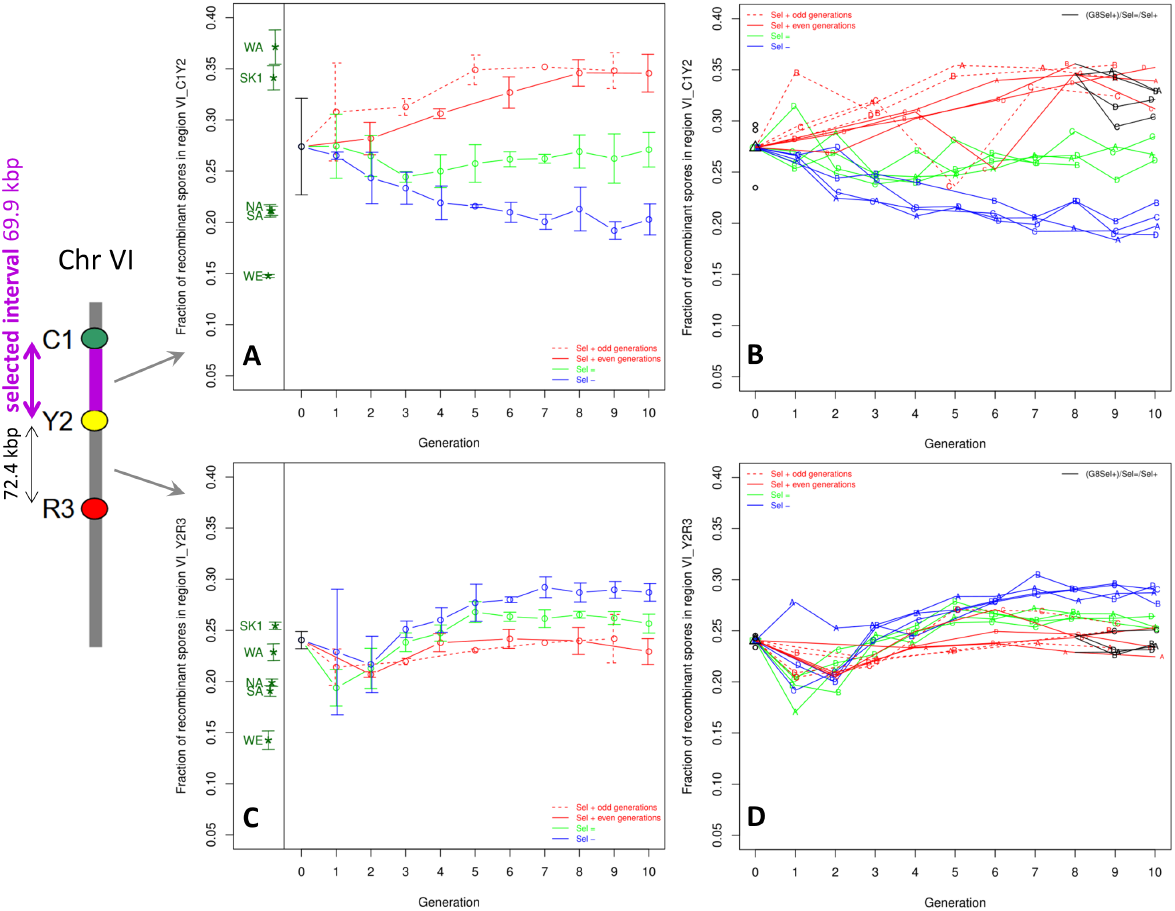
Response of recombination to divergent selection. Fraction of recombinant spores produced by diploids obtained after crossing with a SK1 tester carrying three fluorescent markers delimiting the interval VI_C1Y2, where selection was applied (A, B) and the adjacent interval VI_Y2R3 (C, D). Left panels (A, C): averages and 95% confidence intervals of RR based on four biological replicates. Right panels (B, D): values of each individual replicate (named with colored letters A, B, C, and D on the plots). Black triangles represent the mean of four technical replicates (black circles) of phenotyping the G0 population. Dashed red lines indicate odd generations of Sel+ experiments. Red, green, and blue solid lines indicate Sel+ (even generations), Sel=, and Sel-experiments, respectively. Black lines in panels B and D correspond to aliquots of generation G8 Sel+ which were submitted to one generation of relaxed selection as for Sel=, and then again to the Sel+ regime. Points and confidence intervals in dark green at the left sub-panels of graphs A and C indicate the values of the five founder parental strains from Raffoux *et al*., (2018a). The chromosome sketch on the left side of the figure indicates the positions of the two intervals considered on chromosome VI and their physical length on the SK1 genome.

We observed higher RR values measured in odd than in even generations before G8 (Figure 2A, B), consistent with the additional selection applied to odd generations because of our phenotyping method based on sorting non-fluorescent spores (see Materials and Methods). While RR changes were largely consistent amongst the four biological replicates (Figure 2B), one exception can be seen at G5 and G6 for biological replicate C, Sel+. This is justified a posteriori since there was a failure of the FACS when sorting spores produced by G4 Sel+ C diploids, leading to no selection on recombination. This affected selection both from G4 Sel+ C to G5 Sel+ C but also from G5 Sel+ C to G6 Sel+ C (see Figure 1A). As a result, further generations (G7 to G10) of replicate C missed two generations of efficient selection, and thus, replicate C was removed from G5 to G10 in the calculated means (Figure 2A, C).

When looking at RRs in the immediately adjacent interval Y2R3 on chromosome VI (Figure 2C), RR differed significantly among G0, G10 Sel+, G10 Sel=, and G10 Sel-populations (*p*-value=3.0⨯10^−5^), with pairwise comparisons significant for G0 vs G10 Sel-(*p*-value=9.3⨯10^−5^), G10 Sel+ vs G10 Sel= (*p*-value=0.010), G10 Sel= vs G10 Sel-(*p*-value=0.0018), and G10 Sel+ vs G10 Sel-(*p*-value=4.8⨯10^−5^), and non-significant for G0 vs G10 Sel+ (*p*-value=0.34) and G0 vs G10 Sel= (*p*-value=0.080). However, the responses were in opposite directions as compared to what happened in the VI_C1Y2 interval, where selection was applied. Specifically, when looking at the evolution of RR across generations (Figure 2C), the curves and their confidence intervals suggest that between G0 and G4 or G5, RR in Sel+ became lower than in Sel=, and RR in Sel-became higher than in Sel=. In further generations, RR seemed to reach a plateau during which the values of Sel+ and Sel-relative to Sel= were approximately −12% and +13%, respectively. RR data for all individual replicates are available as Figure 2D, and do not show any particular difference across biological replicates of a given selection mode.

A number of different processes can drive changes in CO numbers, including altered numbers of DSBs, modification of CO interference intensity that typically modulates the CO/NCO ratio, or changes in other steps of meiosis (see review across various animal species in Szasz-Green *et al*. 2025). To get insights on this question, we measured CO interference in our evolution lines. Interference estimated via the coefficient of coincidence (CoC) was significantly positive (CoC<1) in most generations and did not show any pairwise significant difference among G10 Sel+, G10 Sel-, and G10 Sel= populations (*p*-values>0.16) in the region including the selected interval (Suppl Figure S7A, B) and the immediately adjacent one (Suppl Figure S7C, D).

### 2. Divergent selection reshapes the distributions of RR values

While the selection process seemed to successfully generate high- and low-recombining populations on average, it was unclear how penetrant these traits were on a population level. Therefore, to gain insights into the distributions of RR values within each of the evolved populations, we FACS-sorted 1,920 isolated bi-fluorescent diploid cells from generation G8 and phenotyped each of the derived lineages for recombination. Because the FACS cannot distinguish between diploid cells homozygous or hemizygous for the fluorescent markers, only part of these strains were hemizygous, and thus suitable for RR measurement (see Materials and Methods and Suppl Figure S3). Among them, we measured RR in 293 Sel+, 101 Sel=, and 376 Sel-strains. The other strains were discarded.

Average RR values for each selection mode were 0.29 for Sel+, 0.26 for Sel=, and 0.19 for Sel-(Figure 3A), and these three values were significantly different pairwise (Bonferroni-corrected Wilcoxon rank sum tests, *p*-values<0.00018). Average RR values were not significantly different across biological replicates for Sel= strains (adjusted *p*-value=1). However, in Sel+ strains, average RR values differed significantly (*p*-value=0.047) between replicate C (RR=0.27) and replicates A (RR=0.30), which may be related to the different selection regimes applied to replicate C at generations G5 and G6 (Suppl Table S3; Suppl Figure S8).

**Figure 3:**
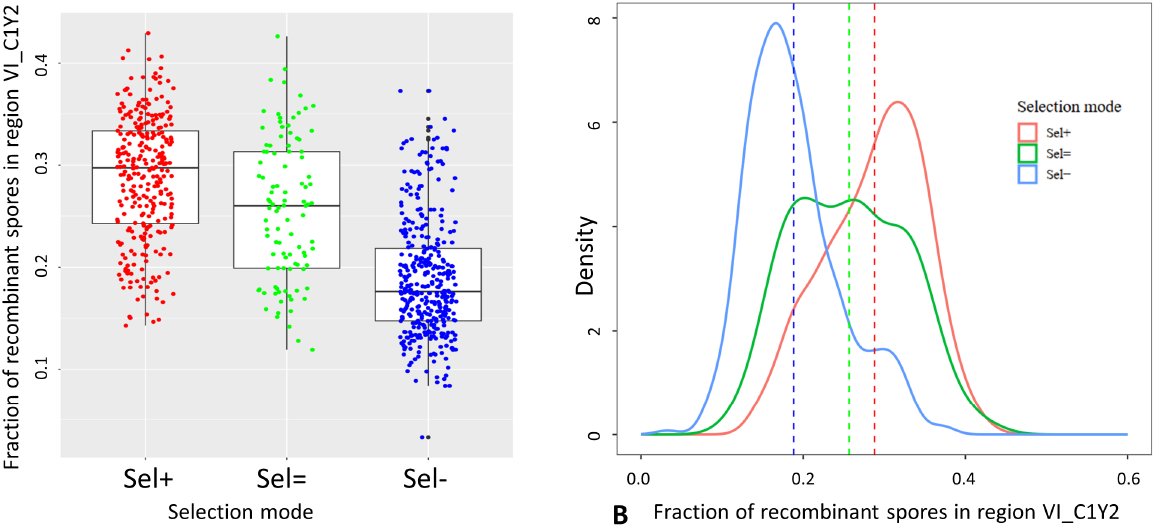
Recombination rate of individuals isolated from G8 populations. Individual values of the fraction of recombinant spores produced by diploids hemizygous for fluorescent markers delimiting the interval VI_C1Y2 (A) and their smoothed distributions (B) measured from 323 Sel+, 377 Sel-, and 101 Sel= isolated from populations at generation G8, after pooling biological replicates A, B, C, D. Dashed lines indicate the means of distributions.

Figure 3B shows the smoothed distributions of RR values for each of the three selection modes. The Sel= distribution was approximately symmetrical (skewness γ1=0.144, MGG symmetry test, *p*-value=0.512), whereas the Sel+ distribution was significantly negatively skewed (γ1= −0.317, *p*-value<2.2⨯10^−16^) and the Sel-distribution was significantly positively skewed (γ1=0.751, *p*-value<2.2⨯10^−16^). The variance of RR in Sel-strains was significantly lower (Fligner-Killeen test) than in Sel= strains (*p*-value=0.0008) and also lower than in Sel+ strains, although less clearly so (*p*-value=0.047). The variance of Sel+ strains showed a tendency to be lower than in Sel= strains, though not significantly (*p*-value=0.052). The overall shapes of these distributions were significantly different for all pairs (KS tests, *p*-values<0.00085). This suggests that selection successfully enriched the initial population for individuals carrying high- and low-recombining genotypes.

When isolating bi-fluorescent diploid cells from G8 Sel+ populations, we found two individuals with outlier RR close to 0.5. Such apparent independent segregation of two linked markers may be due to either a drastic increase in RR, or a change in the chromosomal configuration of the fluorescent markers, for instance, a translocation or duplication of one marker, or a drastic change in the physical length of the interval between the markers. To investigate these possibilities, we whole-genome sequenced one of these two diploid strains. The results showed that the two markers are still at the predicted positions on chromosome VI, that neither of the markers are duplicated at another genomic location, and that both markers are at the hemizygous state (see Suppl Text section 8 and Suppl Figure S9). Moreover, Mendelian segregation of both markers observed by flow cytometry analyses confirmed that they are each present at a single locus. Finally, the sequence between the two markers (85 kbp) in the sequenced strain was homozygous and almost identical to the SK1 sequence, ruling out the idea that some large DNA fragment could have been inserted between the markers. We therefore conclude that the individual has truly attained a high RR.

### 3. Selection for high recombination acts on both local (*cis*) and genome-wide (*trans*) levels

Following ten generations of divergent selection, RR responded to selection within the selected interval and in opposite directions in the immediately adjacent one (Figure 2A, 2C). However, there appeared to be no differences in RR between Sel+, Sel- and Sel= in other probed regions of chromosomes I and VI (Suppl Figure S10, Suppl Figure S11). There was also no significant CoC difference among selection modes at G10 in other probed regions of chromosomes I and VI (Suppl Figure S7 C, D, E, F), but CoC values at G0 were significantly higher than at G10 in both chromosome VI regions (*p*-value<0.0044).

Collectively, these results suggest that the observed response to selection involved primarily local (*cis*) mechanisms. However, the fluorescent markers used in this phenotyping only probe a very limited proportion of the genome and therefore do not accurately reflect genome-wide recombination patterns, nor do they provide insight into features that may have been selected on the local (*cis*) level. To gain a more complete view of genome-wide RRs, as well as of local (*cis*) factors that may have been selected for, segregants of individual meioses were whole-genome sequenced. Given the asymmetric distribution of recombination rates within the Sel+ experiments (see before, and Figure 3B), we reasoned that this population may contain lineages with truly high recombination rates as well as some passenger lineages with near wild-type recombination rates. We therefore sequenced the eleven highest-recombining [high] and the eleven lowest-recombining [low] individuals, all isolated from the Sel+ populations at generation G8 (two to three individuals from each biological replicate A, B, C, and D).

Briefly, tetrads from each of the 20 individuals were dissected to acquire four monosporic segregants, which were then sequenced using Illumina short read sequencing. Segregants were sequenced with a minimum read depth of 39X and a mean read depth of 87X (Suppl Figure S12A). No chromosomal aneuploidies or large copy number variations were apparent among segregants (visual inspection; Suppl Figure S13). Using a custom pipeline (further described in Materials & Methods in section ‘Recombination analysis from tetrad whole-genome sequencing’), the genotype compositions of each segregant were reconstructed. It should be noted that the genotype compositions can only be assigned to one of the four parental genotypes within syntenic regions, so this analysis did not reconstruct genotypes within subtelomeric regions. In addition, the WA and the SK1 tester strain genotypes are so genetically close that they were merged into a single genotype for this analysis.

We first wanted to determine whether there had been any selection for pro-recombinergic features on the local (*cis*) level. Strikingly, the WA/SK1 genotypes were heavily enriched within the Chromosome VI selection interval in [high], but not in [low], segregants (Figure 4A; individual tetrads shown in Suppl Figure S12B). This may be because relative to the SK1 tester strain, the parental strain WE has a truncated chromosome VI arm, and the parental strain SA has a deletion within the selection interval (Suppl Figure S12C). Together, these results suggest that there has indeed been a selection for pro-recombinergic genotype compositions on the *cis* level. Namely, by enriching for the WA/SK1 genotype within the selection interval of Sel+ populations, the [high] evolved lines have both maximized structural similarity and minimized nucleotide heterozygosity to the SK1 tester strain.

**Figure 4:**
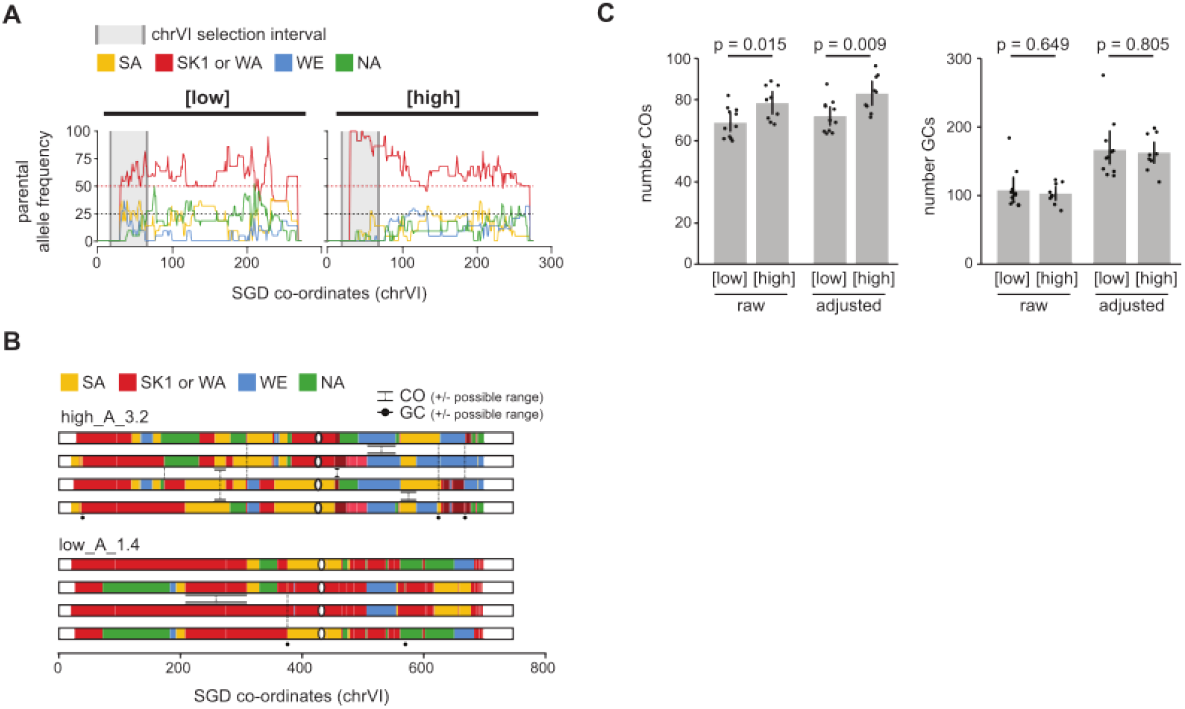
Genome-wide analysis of recombination at G8. (A): Parental allele frequency across chrVI in segregants derived from 9 highest-recombining ([high]) tetrads and 11 lowest-recombining tetrads ([low]) at generation G8 of Sel+ selection. The selection interval used for recombination phenotyping, as well as tetrad selection, is indicated in grey. (B): Examples of possible crossover (CO) and gene conversion (GC) calling in a [high] and [low] tetrad. Given the presence of genetically identical patches within each tetrad, both COs and GCs are under-represented, and precise locations cannot always be determined. (C): Genome-wide quantification of COs and GCs in [low] and [high] segregants, both with and without adjusting for genetically identical patches.

The reconstructed genotypes were next used to determine genome-wide recombination rates. Crossover (CO) events were manually called based on genotype patterns, whereas gene conversion (GC) events were calculated by counting the number of contiguous genome regions containing 3:1 marker patterns (examples shown in Figure 4B).

When quantifying the number of recombination events, it should be noted that because there are ‘patches’ where all four chromatids are genetically identical, the precise location or the precise number of COs within these regions cannot be determined, and GCs cannot be detected at all. Therefore, while some COs occurring within these regions can be inferred based on the surrounding genotype pattern, the total CO number will almost always be under-estimated. This is unlikely to confound our analysis, as the number and cumulative size of such genetically identical blocks are very similar between [low] and [high] populations (Suppl Figure S12D). [high] samples had a slight but non-significant (*p*-value=0.052) increase in the cumulative size of blocks, meaning that CO and GC estimates in [high] samples may be more under-represented than in [low] samples, which is conservative.

To reduce the impact of these genetically identical patches as a potential confounder, CO and GC rate estimates were partially adjusted for each sample according to the size and location of the genetically identical patches. For CO counts, we calculated the size of each genome in which COs could be detected (i.e., excluding any regions that do not have surrounding markers) and normalized the CO counts accordingly (‘adjusted number COs’ in Figure 4C). For NCO counts, we calculated the size of the genome in which NCOs could be detected (i.e., excluding any genetically identical patches) and normalized the NCO counts accordingly (‘adjusted number NCOs’ in Figure 4C). However, it should be noted that this does not fully correct for the under-representation in CO counts, and our detected RR remains lower than expected rates based on existing literature (~90.5 COs per tetrad, Mancera *et al*. 2008). This is because genetically identical blocks that contain surrounding markers might still be large enough to contain multiples of COs, of which only a fraction can be detected with this method. As such, the COs counts depicted here will always be an under-representation of the true CO rates, albeit consistently under-represented between [high] and [low] samples.

Genome-wide, CO rates, but not GC rates, were significantly higher in [high] individuals than in [low] individuals, regardless of whether variations in the genetically identical patches were adjusted for (Figure 4C). Given that COs are relatively rare occurrences and therefore variable between samples, we lack statistical power with our 9 [high] and 11 [low] tetrads to detect significant changes on a chromosomal level. However, it is clear that the increase in CO rate genome-wide is not driven by changes only within the selection interval, with small (non-significant) trends for higher CO rates across multiple chromosomes (Suppl Figure S12F), suggesting that in addition to selecting for pro-recombinergic *cis* elements, at least some [high] lineages have evolved *trans* effectors increasing CO rate remotely. Interestingly, different biological replicates seemed to have different genome-wide responses to selection (Suppl Figure S12E). Biological replicates A and B showed a more consistent increase in global COs following selection, while biological replicates C and D showed either no response (C) or a milder response (D) to selection.

### 4. Single-generation response of RR when relaxing and re-applying selection

Due to a FACS failure, the biological replicate C of Sel+ selection was not properly sorted from G4 to G5, where actually Sel= was applied instead of Sel+. Figure 2B shows that RR dropped down to the Sel= level in that replicate C during the single G4 to G5 transition. Proper Sel+ was then applied from G5 to G6, but given the fact that all G4 spores were kept for G5 in the replicate C, the next spore selection from G5 to G6 (i.e., selecting bi-fluorescent and non-fluorescent spores, see Figure 1A, C) did not, in fact, particularly select recombinant spores. Thus, as expected, RR in G6 Sel+ C stayed at the same level as in G5 Sel+ C. Selection was efficient at the next generation when sorting from G6 to G7 Sel+ C. Consistently, RR went up from G6 to G7 during that single generation, interestingly up to a level higher than in G4.

To try to reproduce such a rapid change in RR and further investigate possible different evolutionary trajectories among biological replicates, we took aliquots of the four biological replicates of G8+, and for each of them, we carried out one generation of Sel= followed by one generation of Sel+ (black lines in Figure 2B). For replicates A and B, RR did not change much during those two new generations, whereas for replicates C and D, RR values tended to decrease. These replicates C and D thus partially reproduced the rapid drop in RR that was observed in G5 Sel+ C and G6 Sel+ C, although RR values did not reach the level of Sel= populations as in G5 Sel+ C and G6 Sel+ C.

To investigate whether some non-genetic mechanisms might have played a role in the quick responses observed previously when relaxing and re-applying positive selection in replicate Sel+ C, we carried out two generations of Sel+ and Sel= as before, but instead of a genetically diverse G0 population, we started from a diploid SK1 strain without any genetic diversity except the hemizygosity of the two fluorescent markers in the same configuration as in the previous G0. From a strict genetic point of view, no response to selection should be expected then. And in fact, we did not observe any difference between Sel= and Sel+ evolutions for RR, either in the interval selected (Suppl Figure S14A), the immediately adjacent interval (Suppl Figure S14B), or the other regions on chromosomes I and VI (Suppl Figure S14D, E, G, H). Similarly, CoC did not differ between Sel= and Sel+ in all three regions (Suppl Figure S14C, F, I).

### 5. Fitness effects of divergent selection on RR

To investigate possible variations of fitness as an associated response to divergent selection on recombination, we first assessed the extent to which the presence and/or the expression of fluorescent markers may introduce confounding effects on fitness. To do so, we sorted spores from G0 having four different configurations of fluorescent markers [++], [+Y], [C+], and [CY], and measured fitness components on the resulting diploids after panmixis within each category. We observed no significant effect of the fluorescent marker status on growth kinetics parameters maximum rate (*p*-value=0.93; Suppl Figure S15A) and plateau (*p*-value=0.56; Suppl Figure S15B). However, a significant effect was detected on sporulation rate (*p*-value=0.035) with [+Y] diploids sporulating significantly better than [C+] ones (*p*-value=0.033; Suppl Figure S15C), and on spores viability (*p*-value=0.0040) with [+Y] spores being significantly more viable than [C+] ones (*p*-value=0.0021, Suppl Figure S15D). Two-factors analysis of variance indicated that the presence of the Y marker insertion had a significant positive effect on spore viability (*p*-value=0.022) and that the presence of the C marker insertion had a significant negative effect (*p*-value=0.0047), but that there was no significant interaction between their effects, so the positive effect of Y might be counteracted by the negative effect of C in [CY] cells.

Then, to assess the fitness of evolved populations, we sorted non-fluorescent spores from populations G0, G8 Sel+, G8 Sel-, and G8 Sel=, obtained diploids from each category by panmixis, and measured fitness components as before. We observed no significant difference between the four populations for growth kinetics parameters maximum rate and plateau (Figure 5A, B), as well as for sporulation rate (Figure 5C). However, spore viability in G8 Sel+ populations was significantly higher than in G0 (*p*-value<10^−4^) and in G8 Sel= (*p*-value=0.023) populations, and spore viability in G8 Sel-populations was significantly higher than in G0 (*p*-value<10^−4^) and in G8 Sel= (*p*-value=0.017) populations (Figure 5D).

**Figure 5:**
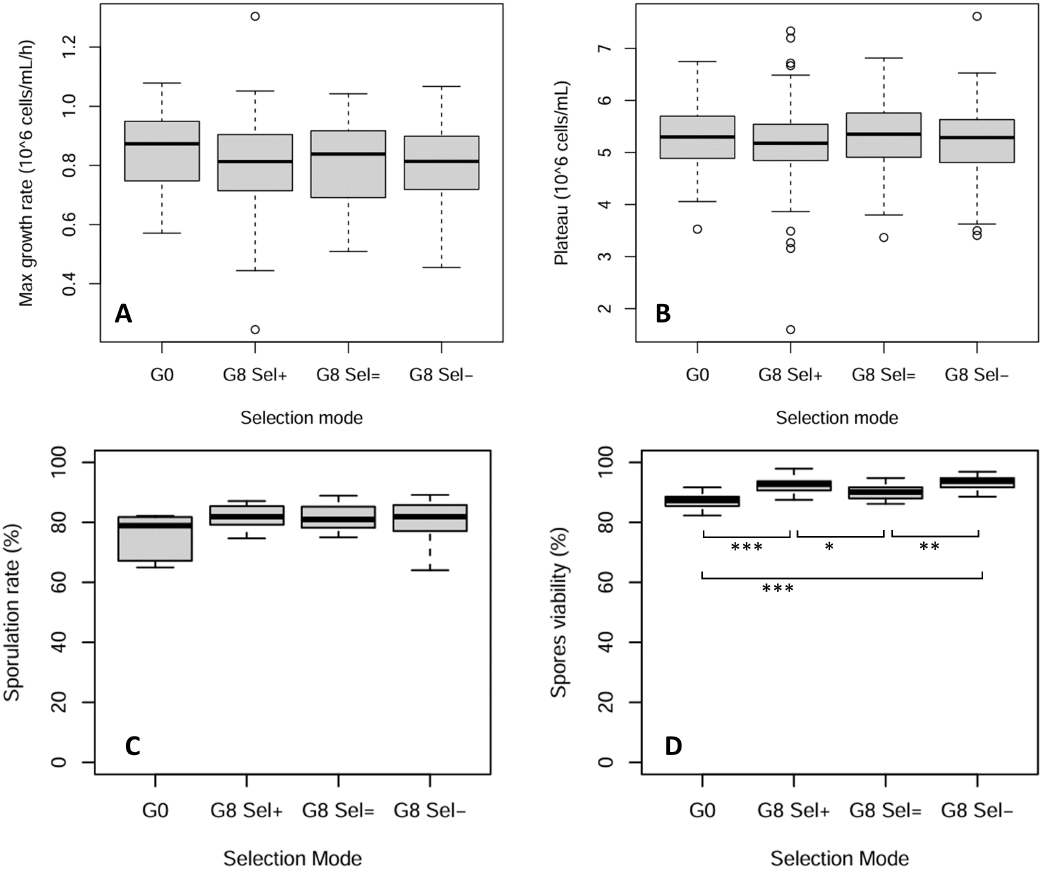
Effect of selection for recombination on fitness. Maximum growth rate (A), concentration at the plateau (B), sporulation rate (C), and spore viability (D) of populations at generation G8 for Sel+, Sel=, and Sel-selection modes, and of the initial population G0. The four biological replicates were pooled. Boxplots were made from a minimum of 30 independent measurements for A and B, and of 6 (C) or 8 (D) independent countings of 96 spores. Pairwise statistical comparisons shown on the graph correspond to Bonferroni-adjusted *p*-values. **<0.05, <0.01**, ** <0.001.

## Discussion

The present divergent selection experiment showed a response of local recombination rate after ten generations of direct selection of recombinant or non-recombinant spores in a 69.9 kbp region of chromosome VI. This region was chosen among the fluorescently-marked intervals previously constructed in the lab, based on the highest variance of RR across the five founder strains (Raffoux *et al*., 2018a), in order to maximize the chance of having alleles of opposing effects segregating for recombination modifiers in our populations. The experiment was designed to investigate evolutionary mechanisms during a limited number of generations. The response to selection we observe is thus most likely due to selection and recombination among allelic variants present in the starting population, rather than accumulated novel mutations. To further limit the occurrence of mutations, we kept the number of mitotic cell divisions low (approximately 10 mitotic cycles per sexual generation). On the other hand, to maximize the standing genetic variation, four of the five founder parents used to build the initial SGRP-4X population were chosen to capture as much as possible of the genetic diversity of the *S. cerevisiae* species (Cubillos *et al*. 2013). Indeed, we observed a plateau in the response to selection after about eight generations (Figure 2A), suggesting that we’ve reached the limits allowed by the initial genetic variation and thus that the impact of novel mutations on that response was probably limited.

The experimental protocol used in this work was designed to sort a total of 300,000 spores at each generation to contribute to the diploids for the next generation, whatever selection mode was applied (Sel-, Sel=, or Sel+). Thus a comparison of RR results across all selection modes is justified in spite of differences in the sorting regimes applied (see Figure 1). The main bottleneck in our experiments is due to FACS, so population size should never be less than 300,000 throughout all steps of all sexual generations. Moreover, spores were efficiently selected from diploids at each cycle as described in Raffoux *et al*., (2018a), avoiding the accumulation of haploid or diploid vegetative cells over generations, which could generate biases and population bottlenecks (Burke, 2023). Genetic differentiation between G0 and Sel= populations is thus expected to be mainly due to unwanted selection (e.g. on growth, sporulation, viability) rather than genetic drift. Therefore, using Sel= populations as a control should allow us to investigate the effect of selection on recombination without confounding effects of drift or unwanted selection on other traits.

We do not know of any similar experiment performed previously in *S. cerevisiae*. In S. pombe, an experiment of evolution under selection for recombinant or non-recombinant spores was recently reported in a PhD dissertation (Berenguer Millanes 2025). The author carried out a similar approach based on FACS-sorting spores expressing fluorescent proteins, but the initial diploid was obtained by crossing two quasi-isogenic variants (differing only at a few marker loci) from a single Leupold lab strain, so the main source of genetic variance allowing for evolution was novel mutation. After 36 meiotic generations, the author observed a significant positive response when selecting double-recombinant spores in two adjacent intervals on Chromosome I, but single interval results were unclear. It should be noted that (1) in such selection for double recombinants, the effective selection pressure applied on RR is much stronger than in our case (single interval) because it requires two different COs, and (2) the absence of crossover interference in S. pombe also makes simultaneous selection on two close intervals more effective than in *S. cerevisiae*. More generally, quantitative genetics studies within and between animal species suggest that genome-wide RR is expected to display a weak response to short-term selection (Payseur 2025). This large compilation of results sheds light on the evolution of global *trans*-acting factors modifying RR in nature, but its conclusions are not necessarily expected to apply to our present study, which involves laboratory selection on COs in a specific region, with an artificial population designed to maximize its initial genetic variation. In our case, the experimental design was more prone to revealing *cis* sequences modulating RR locally, for instance via the level of heterozygosity. Our results are consistent with previous works showing that RR is able to evolve under selective pressure, e.g., in Drosophila (Detlefsen and Roberts 1921; Chinnici 1971a; Chinnici 1971b; Charlesworth and Charlesworth 1985a) and other species, as might be expected given the high extent of variation of RR observed across taxa, see large review in (Stapley *et al*. 2017).

Our evolved populations were initially obtained by crosses between five strains (SK1, WA, NA, SA, and WE), whose RRs have been previously measured (Raffoux *et al*., 2018a) after crossing with the same SK1 fluorescent testers as in the present work. It is thus interesting to compare the RRs between our evolved populations and their parental ancestors, as shown in Figures 2A, 2C, Suppl Figures S10A, S10C, S11A, and S11C. First, the SK1 parent, which corresponds to completely homozygous diploids, generally shows the highest RR values. In the selected interval VI_C1Y2 (Figure 2A), which has been chosen for its high RR variance among the parents, G0 and G10 Sel= RRs roughly lie in the middle of the five parental values, whereas G10 Sel+ RR almost reaches up to SK1 or WA values. Analogously, G10 Sel-RR reaches down to NA and SA values but is still higher than WE. Our recurrent selection thus did not generate transgression, and this is compatible with the hypothesis that Sel+ populations increased their RR mostly by evolving homozygous sequences in the selected interval, thus reaching RR levels comparable to highly homozygous SK1xSK1 or SK1xWA diploids, whereas Sel-populations did not reach homozygosity levels as low as in the WExSK1 hybrid. In the other intervals, our populations generally show RR values higher than the parental average and close to those of the highest parents (mostly SK1). This was expected since SK1 alleles have a theoretical frequency of 50% in our populations, leading to frequencies of homozygous diploids much higher than if all parents contributed equally.

In our experiment, selection was stronger in Sel+ than in Sel-since spores selected in Sel-may result from hidden recombination between the markers (Figure 1A). Quantitatively, the expected frequency of spores produced with a (hidden) recombination event between the marker loci, among all spores kept by the FACS in Sel-regime, ranges between zero and 1/3 as RR ranges between 0 and 1/2, so one could expect selection to be more efficient in Sel+. However, we observed no clear difference in the absolute values of slopes of the responses between Sel+ and Sel-(Figure 2A).

We observed an inverted response to divergent selection in the interval adjacent to the one where selection was applied. This may be due to CO interference, which is significantly positive in the interval VI_C1Y2R3 (CoC<1; Suppl Figure S7A): if local determinants drive double-strand breaks to be preferentially repaired into COs in one region, then interference is expected to commit double-strand breaks surrounding that region to the NCO pathway, thus depleting COs in adjacent intervals. When we assessed the strength of interference along generations, no significant changes were observed between Sel+, Sel-, and Sel= experiments, indicating that changes in interference strength were not the main mechanism that led to modified CO frequency as a response to selection. In the interval VI_R3Y4C5, we measured surprisingly high values of CoC (greater than one in the first three generations) whereas CoC values became stable and lower than 1 in further generations. Values of CoC greater than one are supposed to indicate negative interference, which is not in accordance with our previous measurements of positive interference (corresponding to gamma parameter nu>1) with the same method in the founder strains of the SGRP-4x population (Raffoux *et al*. 2018a). Given the high variance associated with the data, we suggest that negative interference measured in generations G0 to G2 should be taken with care. Nevertheless, in other regions too, we observed a similar tendency of interference to increase (i.e., of CoC to decrease) across generations, whatever the mode of selection, Sel+, Sel-, or Sel=. If this increase is related to genetic changes in the population, it cannot be due to our selection on recombination, but might be a consequence of unwanted selection on some other traits, which is unavoidable in experimental evolution, particularly when sporulation is involved in the selection cycle. Elsewhere, our hypothesis that the changes in RR achieved in our Sel+ populations within the selected interval may be due to a shift in resolution of CO/NCO decisions may suggest that increase in RR would be associated with a decrease in interference, which is not supported by our CoC results. Still, in many results on the determinants of recombination variation in animal species, shifts in the CO/NCO choice was shown to be a major pathway (Szasz-Green *et al*. 2025).

Phenotyping individuals isolated from the evolved populations revealed that eight generations of directional selection reduced the variance of RR values in the selected interval for Sel+ and Sel-experiments, but not in the Sel= control. This may be explained by the fact that strong directional selection in finite populations generally reduces genetic diversity (Walsh and Lynch, 2018), which in turn results in reduced phenotypic variance for that trait. Still, in both Sel+ and Sel-evolved populations, we observed continuous distributions of RR (Figure 3B) with gradual changes in the mean across generations as compared to Sel= (Figure 2A), pointing to the fact that we’ve successfully evolved population-level changes in the recombination rates, without solely selecting for some strong recombination variant. However, the shapes of the distributions were also modified, with a negatively skewed distribution in G8 Sel+ and a positively skewed distribution in G8 Sel-, while the distribution remained symmetrical in G8 Sel= (Figure 3B). These two asymmetric distributions might be explained by considering the hypothesis of a rather strong QTL affecting RR, with two alleles (‘high-RR’ and ‘low-RR’) segregating in our populations. The Sel+ population would mainly contain individuals carrying the ‘high RR’ allele at this QTL, visualized by the highest peak on the right side of the red curve in Figure 3B, and a minority of individuals with the ‘low RR’ allele revealed by the shoulder on the left side of the red curve. Symmetrically, the Sel-population would mainly contain individuals carrying the ‘low RR’ allele at this QTL, visualized by the highest peak on the left side of the blue curve in Figure 3B, and a minority of individuals with the ‘high RR’ allele revealed by the shoulder on the right side of the blue curve. In Sel=, both alleles would have similar frequencies, which would result in the observed symmetrical distribution. Moreover, the Sel= curve shows three peaks, and the shapes of Sel+ and Sel-curves also suggest a possible third bump between the two main parts of the peaks, which might correspond to the three genotypes of a codominant QTL. Elsewhere, in Sel-, the peak is higher than in Sel+, indicating a greater loss of variance. This may suggest that selection was stronger in Sel-than in Sel+ experiments, which is not expected given our experimental design allowing some hidden recombinants to be selected in Sel-. Another hypothesis is that higher recombination in Sel+ increased the genetic variance by creating new allele combinations, thus reducing the loss of variance due to selection.

Following selection within the chromosome VI_C1Y2 interval, the responses in RR in the five other intervals carrying fluorescent markers suggest a *cis*-only change in RR. However, the regions probed by fluorescent markers are very limited as compared to the whole genome, leading us to sequence individual [high] and [low] segregants from G8 Sel+ to obtain a genome-wide and higher-resolution picture of recombination landscapes. When looking at the parental allele frequencies within the selection interval, we found that [high] samples almost always showed an enrichment for the WA/SK1 genotype, suggesting a strong selection on the *cis* level. This is perhaps unsurprising given the large structural variants of the other parents within the selection interval, which might hinder efficient homolog pairing and subsequent recombination (Suppl Figure S12). However, it should also be noted that SK1 has been widely used in meiotic recombination studies due to its pro-recombinergic features (e.g., efficient sporulation). It is therefore possible that the local enrichment of WA/SK1 is due to both the sequence-level and structural similarity of WA with the initial SK1 tester strain, or due to a subsequent enrichment of SK1 within the selected interval due to some of its pro-recombinergic features, or a mixture of both. When looking genome-wide, we observed a significantly higher genome-wide CO number in [high] compared to [low] tetrads. Given that COs are relatively rare on a per-chromosome level, the data are noisy and statistical power is low, but we also observed a (non-significant) trend to increase CO number on chromosomes not containing the selection interval. Together, these suggest a possible additional selection on the *trans* level.

Intriguingly, the tetrads sequenced from the four G8 Sel+ evolution lines (i.e., biological replicates A, B, C, and D) appeared to increase the chromosome VI recombination rates through different mechanisms that we were able to discern by sequencing individual segregants from each tetrad. In tetrads from G8 Sel+ A [high] and G8 Sel+ B [high], we observed consistently high genome-wide recombination rates (80, 89, 81, 87, and 87 COs per tetrad), suggesting successful selection for genome-wide increases in recombination rates, as intended. Conversely, in G8 Sel+ C [high] and G8 Sel+ D [high], we did not observe high genome-wide recombination rates (68, 69, 73 COs per tetrad), and we observed no striking abnormalities on the *cis* level. It is therefore unclear why the tetrads sequenced from G8 Sel+ C [high] and G8 Sel+ D [high] display high recombination rates within the chromosome VI selection interval. These different behaviors between replicates A and B on one hand, and C and D on the other hand may be related to the RR changes observed when we relaxed selection for two generations in the four G8 Sel+ biological replicates (black lines in Figure 2B). Such variation in evolutionary outcomes of technically identical experimental setups highlights the power of this approach for uncovering biological novelty.

In the biological replicate C of Sel+ experiment (Figure 2B), when the selection pressure was relaxed (all spores were kept instead of only recombinant ones) between G4 and G5, RR went back to its initial level within just one generation, and recovered its G4 value as soon as recombinant spores were again specifically selected, which occurred only at G7 (see Results, and Figure 1A). When we tried to reproduce the same situation by relaxing spore selection starting from each of the four biological replicates of the G8 Sel+ population, a similar response was observed for replicates A and B, but replicates C and D were not affected by the change in the selection mode (black lines in Figure 2B). These results raise the question of which kind of genetic element or factor affecting recombination could have its frequency massively altered in a single generation without selection, and massively recovered in a single generation when selecting again recombinant spores. Because the action of this element was observed at the local level only, one could think of a sequence motif with major local effect, such as a hotspot, and/or a particular diploid genotype with an inhibitory effect on COs, due, for instance, to heterozygosity of a structural variant such as an inversion or a deletion. Although we have not identified the real mechanism at work in our experiment, it seems to be able to play an important role in the response to directional selection.

Our results show that eight generations of divergent selection on recombination did not change growth kinetics (Figure 5A, B) or sporulation rate (Figure 5C). Given the experimental design involving mitotic growth and sporulation steps, it is likely that some unwanted selective pressure was applied on these traits, so an increase of fitness may have been expected between G0 and G8 generations. However, the SGRP-4X population had already undergone 12 generations of sexual reproduction before we started our experiments (Cubillos *et al*. 2013), so major alleles controlling these fitness components may have already been more or less fixed. On the other hand, spore viability in G8 Sel+ and in G8 Sel-were significantly higher than in G8 Sel= and G0 populations (Figure 5D). Because it increased CO rates to some extent genome-wide, the Sel+ treatment may have reduced the frequency of achiasmatic bivalents leading to aneuploidy, thus increasing the ability of spores to germinate as compared to the Sel= case. However, this does not explain why Sel-populations also gained better spore viability than Sel= ones. A hypothesis might be that this is due to fitness effects of the fluorescent markers inserted in the genome. Indeed, [C+] spores were less viable — but [+Y] spores more viable — than [++] spores (Suppl Figure S15), both effects being more or less additive (see Results). The advantage of being [+Y] might be explained by the fact that severely non-viable spores, which could not express fluorescent proteins, cannot be present in the [+Y] class but may be present in the [++] class, introducing some sampling bias. Besides that, [C+] spores might be less viable than [++] ones because of some deleterious effect of C marker insertion: even though we checked that the insertions did not disrupt any known coding sequence, it might alter some fitness-related regulatory element. Altogether, the gain in viability of Sel-spores as compared to Sel= spores may not be clearly attributed to confounding effects of the fluorescent markers.

In conclusion, the recurrent divergent selection approach used in this work proved to be an efficient way to investigate the ability of the recombination rate to respond to directional selection. A clear response was observed rather symmetrically in both directions in the selected region. *Cis* effects were observed at the sequence level, but *trans* effects were also suggested, providing quite direct evidence that various types of recombination modifiers are at work to control the evolution of recombination through multiple paths. It also suggests that the ones at work in our experiments involve some QTLs with strong effects contributing to the response.

## Data availability statement

Yeast strains produced in this work are available upon reasonable request. All data underlying this article are publicly available in the french government data warehouse ‘Recherche Data Gouv’ (https://recherche.data.gouv.fr/en) under the persistent DOI: https://doi.org/10.57745/YTFYTT. R and Python codes used to analyze the data and produce graphs and conclusions are publicly available in the software forge ‘Forge INRAE’ at https://forge.inrae.fr/gqe-base/evolrec_wp1 which is also mirrored in ‘Software Heritage’ repository (https://www.softwareheritage.org/) under the name ‘evolrec_wp1’ accessible from the permalink https://tinyurl.com/swh-evolrec-wp1.

## Acknowledgements

The authors thank Mickael Bourge for many years of sharing with us his invaluable experience in cytometry analysis, Mickael De Carvalho for having so many discussions with us about the project, Domenica Manicacci for advice on statistical analyses, and two anonymous reviewers as well as the editor for their helpful comments on the manuscript.

## Study funding

The present work has benefited from Imagerie-Gif core facility supported by I’Agence Nationale de la Recherche (FBI ANR-24-INBS-0005 (BIOGEN); SPS ANR-17-EUR-0007, EUR SPS-GSR) and with financial support from ITMO Cancer of Aviesan and INCa on funds administered by Inserm. This work has also benefited from funding from the EVOLREC (ANR-20-CE13-0010) and CO-PATT (ANR-20-CE12-0006) projects. GQE—Le Moulon benefits from the support of the LabEx Saclay Plant Sciences-SPS (ANR-10-LABX-0040-SPS).

## Conflict of interest

The authors declare no conflict of interest

## Notes

https://doi.org/10.57745/YTFYTT

https://tinyurl.com/swh-evolrec-wp1

https://forge.inrae.fr/gqe-base/evolrec_wp1

### Competing Interest Statement

The authors have declared no competing interest.

### Summary of Updates

This version corresponds to the text formally accepted for publication by Genetics and currently in press under the DOI: 10.1093/genetics/iyag239

