## Supplementary material for "Response to divergent selection on meiotic recombination in Saccharomyces cerevisiae": Suppl Figure S1

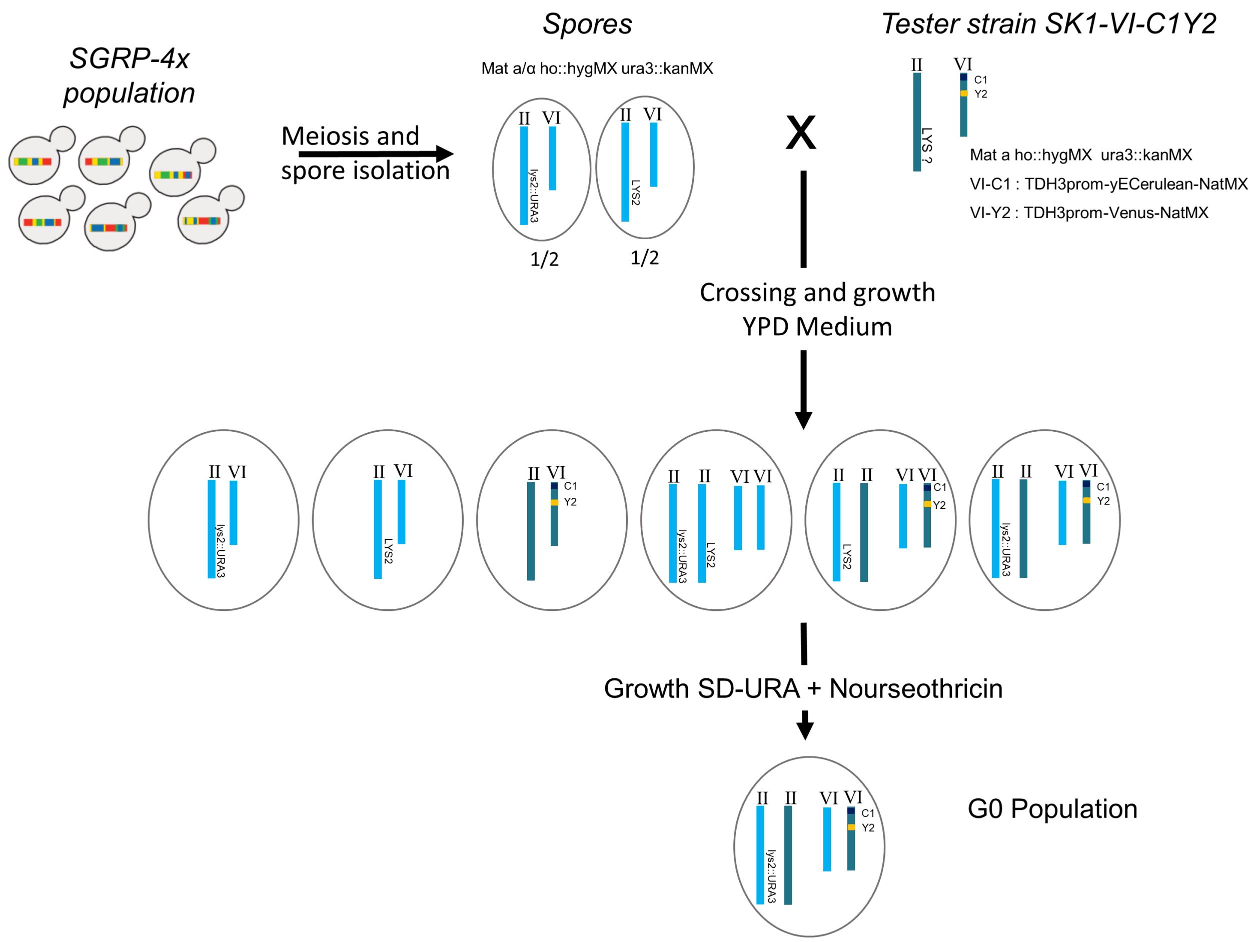


**Suppl Figure S1. Procedure used to build the population G0 from the SGRP-4X population.** Mixed spores from SGRP-4X (Mat-a or Mat-α ho::hygMX ura3::kanMX, lys2::URA3 or LYS2) were mated with a fluorescent SK1-VI_C1Y2 tester (Mat-a ho::hygMX ura3::kanMX VI-C1:TDH3prom-yECerulean-NatMX VI-Y2:TDH3prom-Venus-NatMX). Diploids hemizygous for the fluorescent markers were then selected based on their ability to grow on medium without Uracil containing nourseothricin.
