## Supplementary material for "Response to divergent selection on meiotic recombination in Saccharomyces cerevisiae": Suppl Figure S2

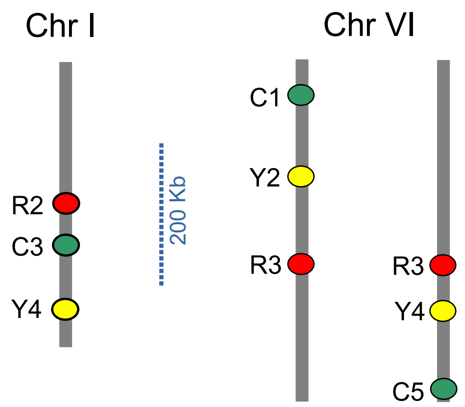


**Suppl Figure S2. Marker positions in the testers used to probe recombination rate.** These testers are SK1 strains containing three fluorescent markers, used to cross to non-fluorescent cells to measure recombination rate in six different intervals: two on chromosome I and four on chromosome VI (among which the VI_C1Y2 interval where selection was applied). C, Y, R refer to markers yECerulean, Venus, and mRFP, respectively.
