## Supplementary material for "Response to divergent selection on meiotic recombination in Saccharomyces cerevisiae": Suppl Figure S3

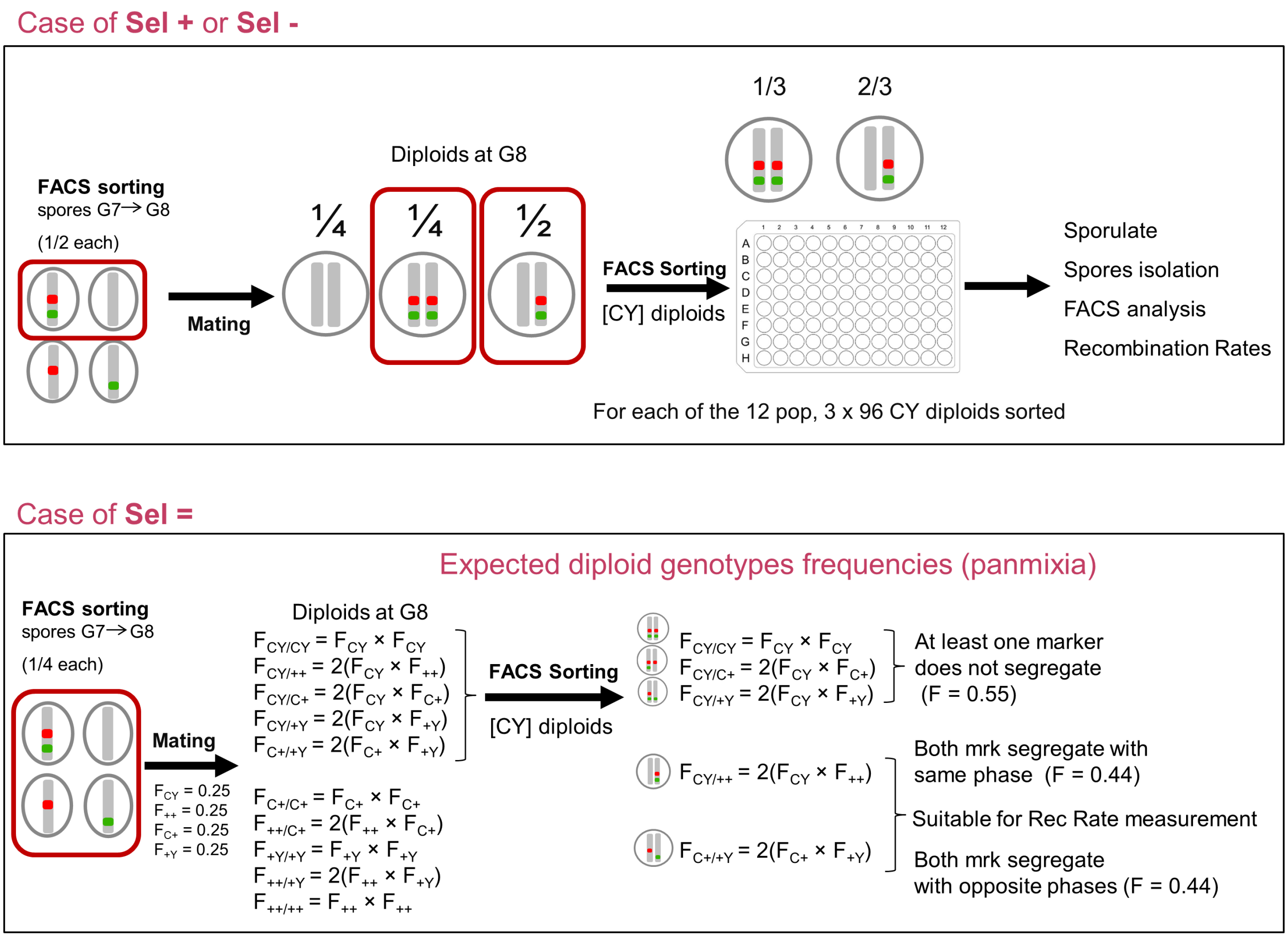


**Suppl Figure S3. Phenotyping recombination rate of individuals isolated from G8 populations**. Experimental design and expected diploid frequencies after FACS-sorting bi-fluorescent diploid cells in the case of Sel+ and Sel- (upper panel) or Sel= (lower panel) experiments.
