## Supplementary material for "Response to divergent selection on meiotic recombination in Saccharomyces cerevisiae": Suppl Figure S4

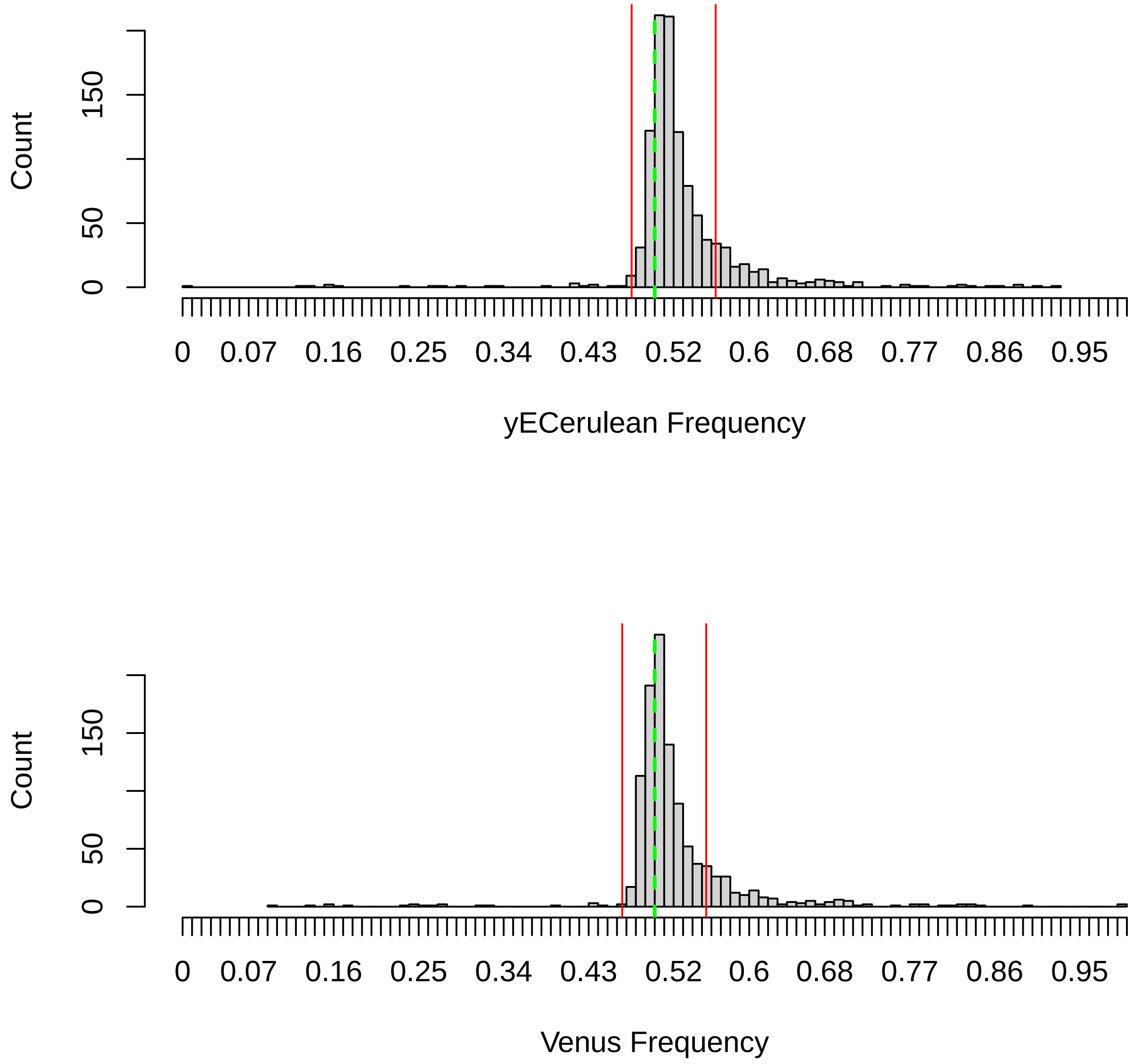


**Suppl Figure S4. Distribution of fluorescent marker frequency in spores from individuals isolated from G8 populations. (A):** Marker yECerulean. **(B):** Marker Venus. X-axis: marker frequency. Y-axis: number of isolated individuals. Red lines: lower and upper boundaries defined as median +/- 3 times Median Absolute Deviation. Green dashed lines correspond to a frequency value of 0.5.
