## Supplementary material for "Response to divergent selection on meiotic recombination in Saccharomyces cerevisiae": Suppl Figure S5

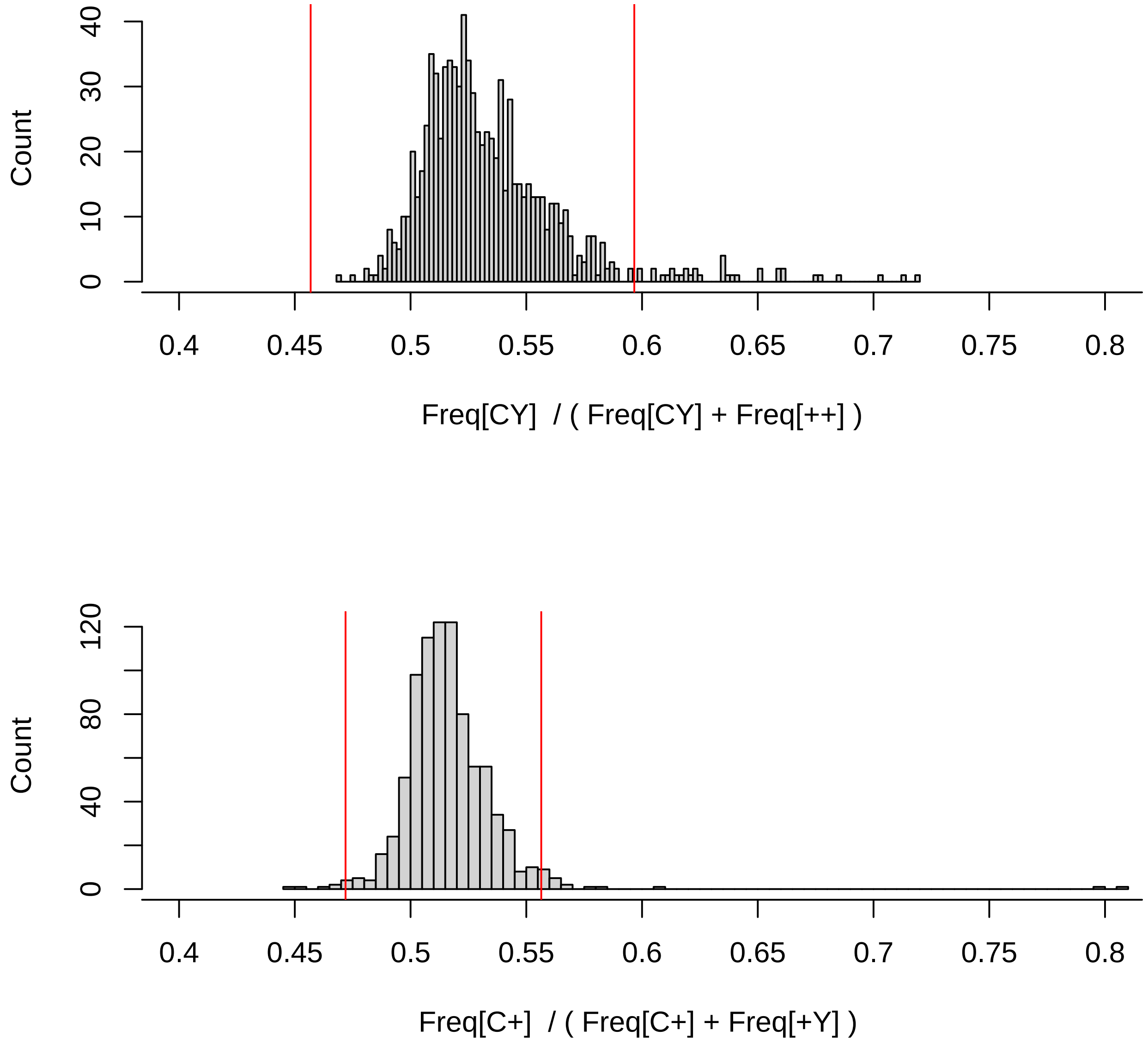


**Suppl Figure S5. Distribution of haplotype frequency ratios in spores from individuals isolated from G8 populations.** X-axis: haplotype frequency ratios $F_{[CY]}/\left(F_{[CY]}+F_{[++]}\right)$ **(A)** and $F_{[\mathrm{C+}]}/\left(F_{[\mathrm{C+}]}+F_{[\mathrm{+Y}]}\right)$ **(B)**. Y-axis: number of isolated individuals. Red lines: lower and upper boundaries defined as median +/- 3 times Median Absolute Deviation.
