## Supplementary material for "Response to divergent selection on meiotic recombination in Saccharomyces cerevisiae": Suppl Figure S6

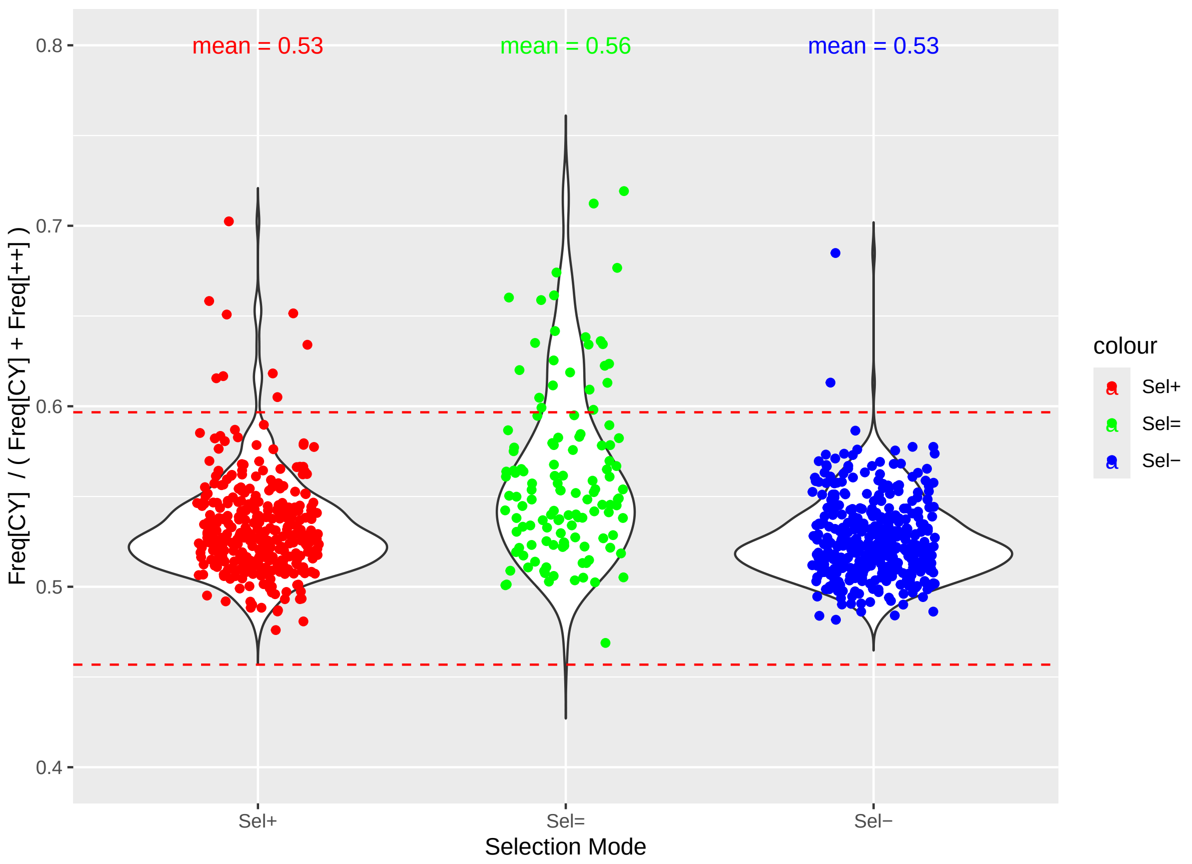
mean = 0.53
mean = 0.56
mean = 0.53
0.8
0.7
0.6
0.5
0.4
Freq[CY] / ( Freq[CY] + Freq[++] )
Sel+
Sel=
Sel−
Selection Mode
colour
Sel+
Sel=
Sel−

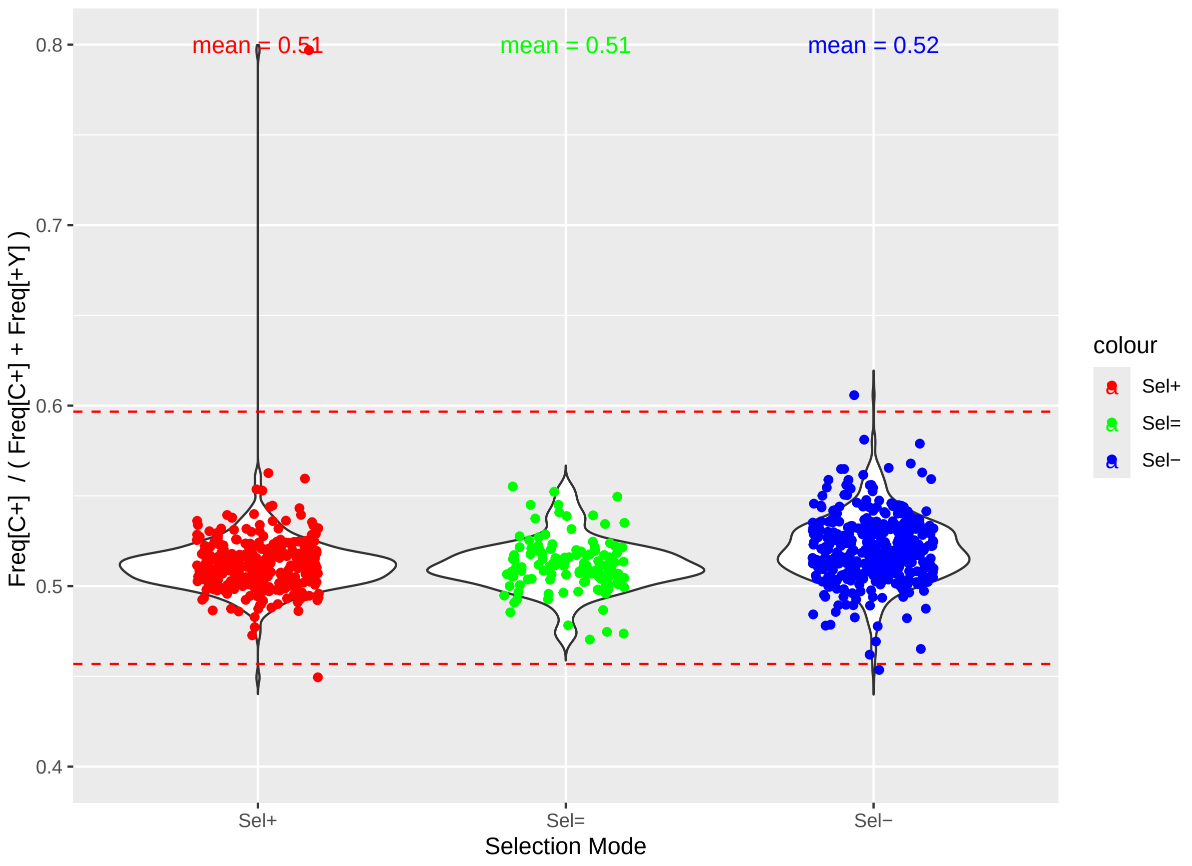
mean = 0.51
mean = 0.51
mean = 0.52
0.8
0.7
0.6
0.5
0.4
Freq[C+] / ( Freq[C+] + Freq[+Y] )
Sel+
Sel=
Sel−
Selection Mode
colour
Sel+
Sel=
Sel−

**Suppl Figure S6. Distribution of haplotype frequency ratios in spores from individuals isolated from G8 populations for each selection mode.** X-axis: selection mode: Sel+ (red), Sel= (green), and Sel- (blue). Y-axis: haplotype frequency ratios $F_{[CY]}/\left(F_{[CY]}+F_{[++]}\right)$ **(A)** and $F_{[\mathrm{C+}]}/\left(F_{[\mathrm{C+}]}+F_{[\mathrm{+Y}]}\right)$ **(B)**. Red dashed lines: lower and upper boundaries defined as median +/- 3 times Median Absolute Deviation.
