## Supplementary material for "Response to divergent selection on meiotic recombination in Saccharomyces cerevisiae": Suppl Figure S7

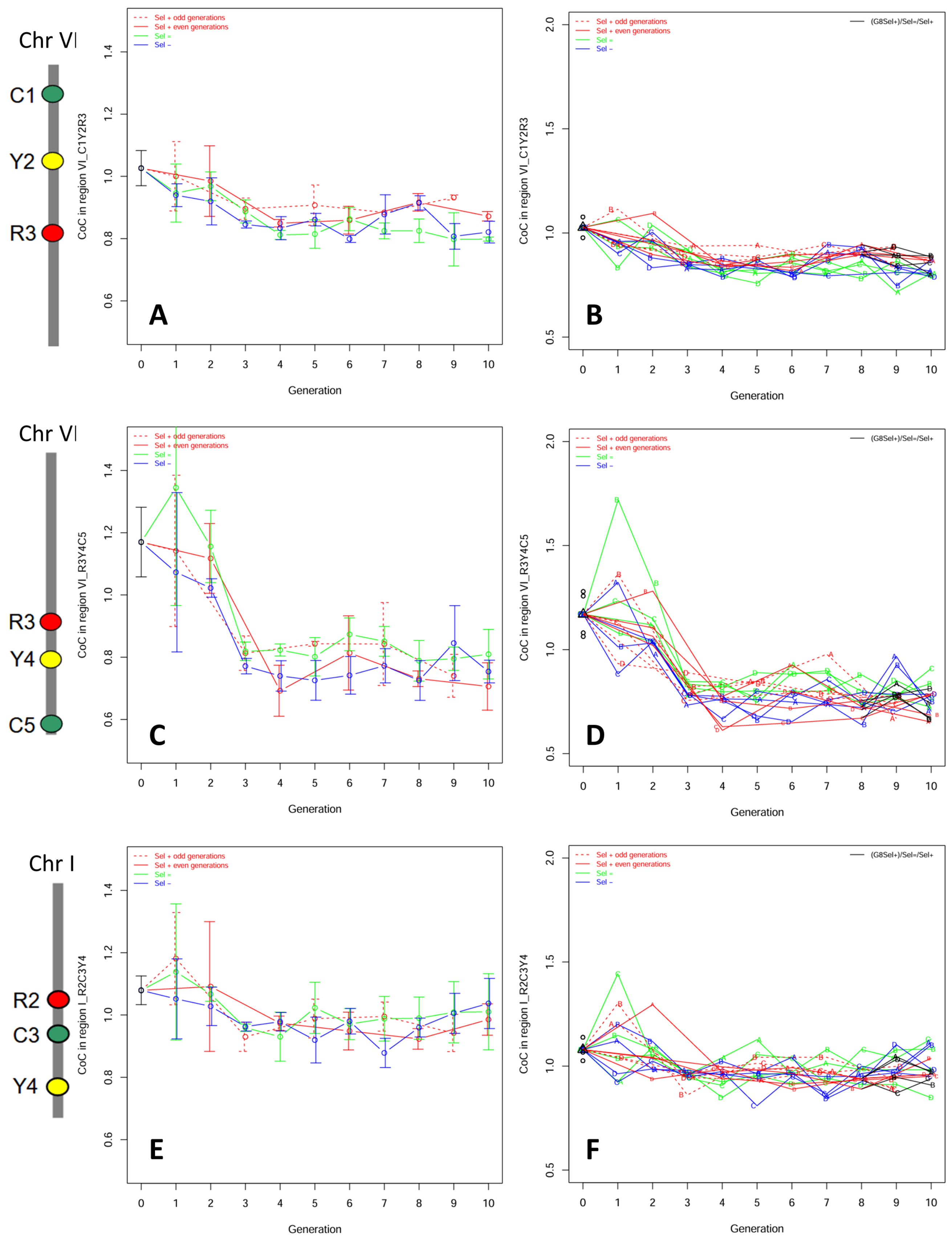


**Suppl Figure S7. Crossover interference across generations of selection.** Coefficient of coincidence (CoC) based on two adjacent intervals delimited by three fluorescent markers, in two regions of chromosome VI **(A, B, C, D)**, and in one region of chromosome I **(E, F)**. CoC=1 corresponds to no interference, whereas CoC<1 corresponds to positive interference. **(A, C, E):**

Mean CoC values and 95% confidence intervals based on four biological replicates. **(B, D, F):** CoC values of each individual biological replicate. Dashed red lines indicate even generations of Sel+ experiments. Red, green, and blue solid lines indicate Sel+ (odd generations), Sel=, and Sel- experiments, respectively. The chromosome sketches on the left side of the figure indicate the positions of the pairs of intervals considered.
