## Supplementary material for "Response to divergent selection on meiotic recombination in Saccharomyces cerevisiae": Suppl Figure S8

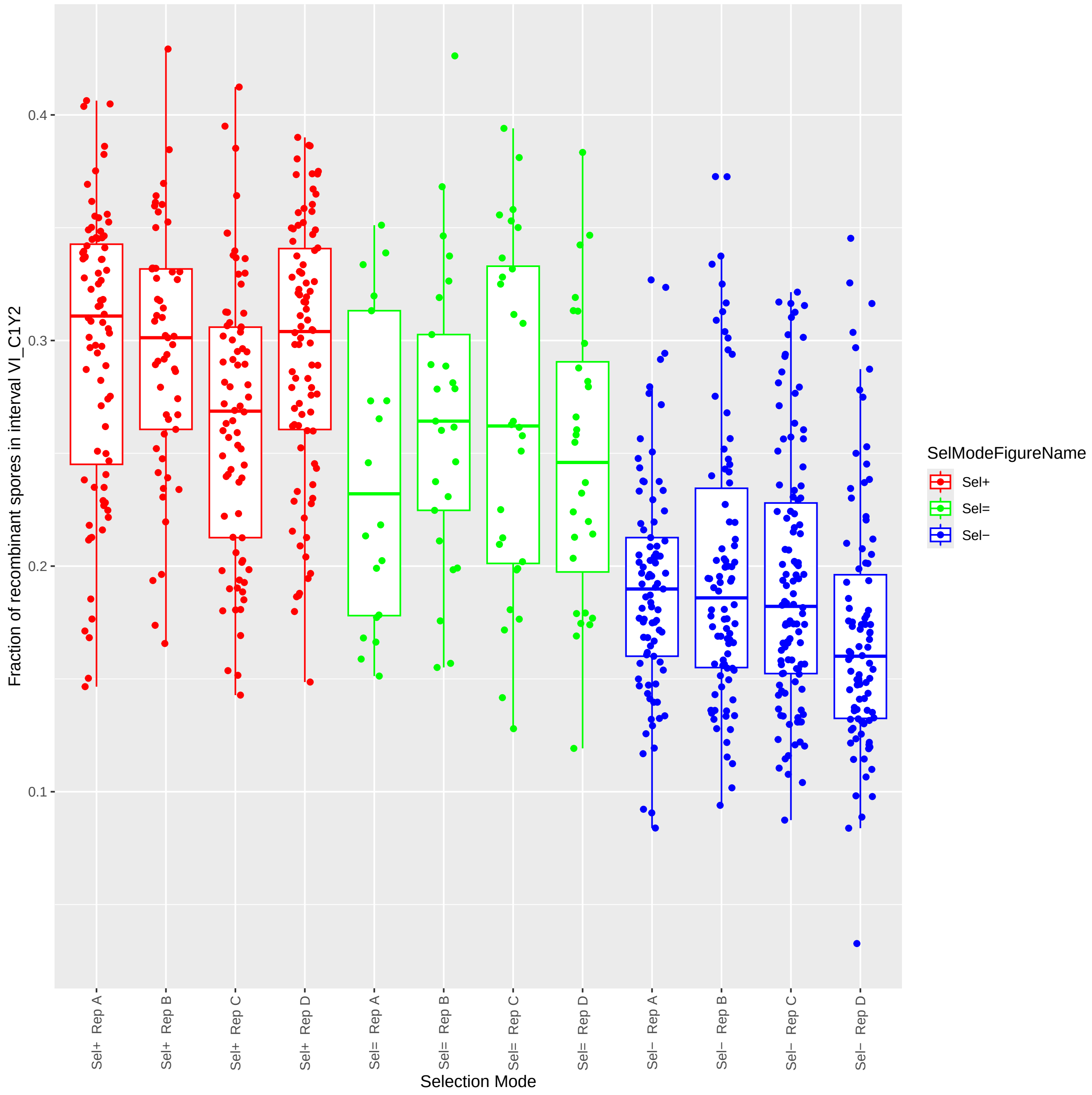


**Suppl Figure S8. Recombination rates of individuals isolated from G8 populations per selection mode and per replicate.** Individual values of the fraction of recombinant spores produced by diploids hemizygous for fluorescent markers delimiting the interval VI_C1Y2 for each biological replicate A, B, C, D, and each selection mode Sel+ (red), Sel= (green), and Sel- (blue).
