## Supplementary material for "Response to divergent selection on meiotic recombination in Saccharomyces cerevisiae": Suppl Figure S9

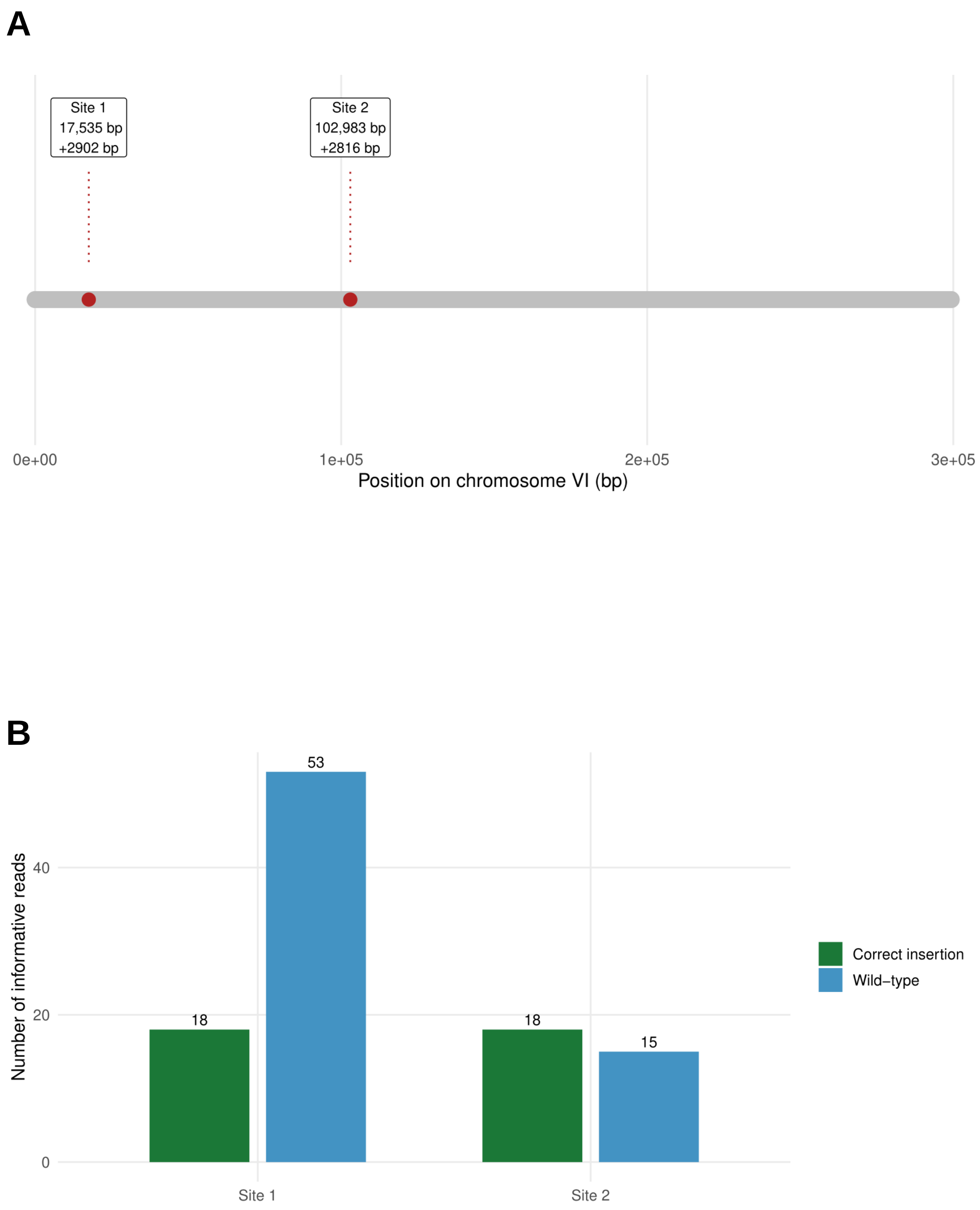


**Suppl Figure S9**. **Sequence analysis of an outlier hyper-recombinant strain. (A)** Map of *S. cerevisiae* SK1 chromosome VI (CP020179.1; 299,318 bp) drawn to scale, showing the two fluorescent cassette integration sites identified from the long-read data; each site is labeled with its break point coordinate and the median across reads of the insertion size. **(B)** Classification of informative reads at each site showing the hemizygous status of the two markers. Green: reads containing the correct-size insertion at the predicted site. Blue; reads without any insertion.
