## Supplementary material for "Response to divergent selection on meiotic recombination in Saccharomyces cerevisiae": Suppl Figure S10

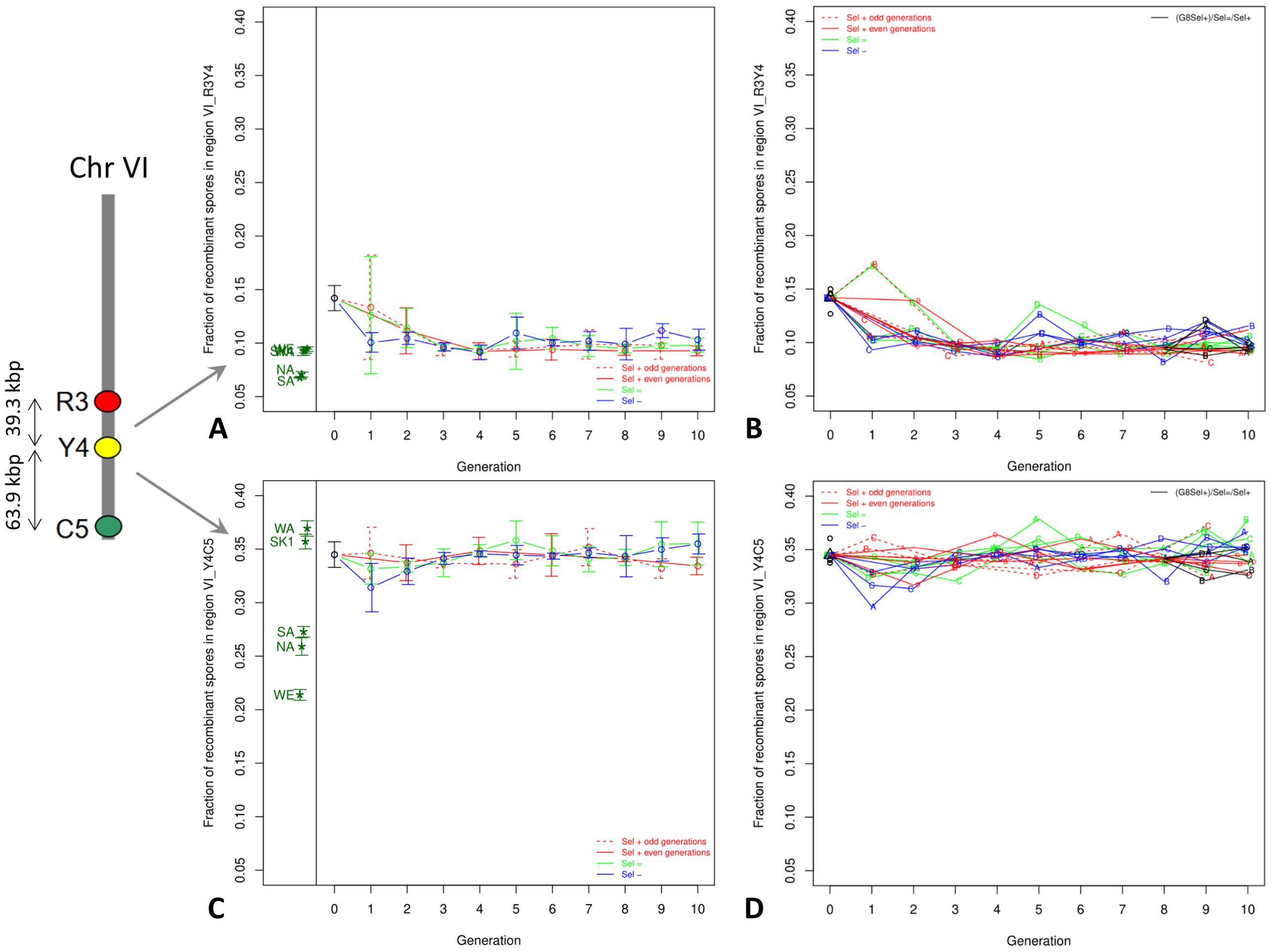


**Suppl Figure S10. Response to selection in two other genomic regions on chromosome VI.** Fraction of recombinant spores produced by diploids obtained after crossing with a SK1 tester carrying three fluorescent markers delimiting the two regions VI_R3Y4 **(A, B)** and VI_Y4C5 **(C, D)**. **(A, C):** Means and 95% confidence intervals based on four biological replicates (named with colored letters A, B, C, and D on the plots). **(B, D):** Individual values of each biological replicate. Dashed red lines indicate even generations of Sel+ experiments. Red, green, and blue solid lines indicate Sel+ (odd generations), Sel=, and Sel- experiments, respectively. Black lines in panels **B** and **D** correspond to aliquots of generation G8 Sel+ which were submitted to one generation of relaxed selection as for Sel=, and then again to the Sel+ regime. Points and confidence intervals in dark green at the left sub-panels of graphs A and C indicate the values of the five founder parental strains from Raffoux *et al.*, (2018a). The chromosome sketch on the left side of the figure indicates the positions of the two intervals considered on chromosome VI and their physical length on the SK1 genome.
