## Supplementary material for "Response to divergent selection on meiotic recombination in Saccharomyces cerevisiae": Suppl Figure S11

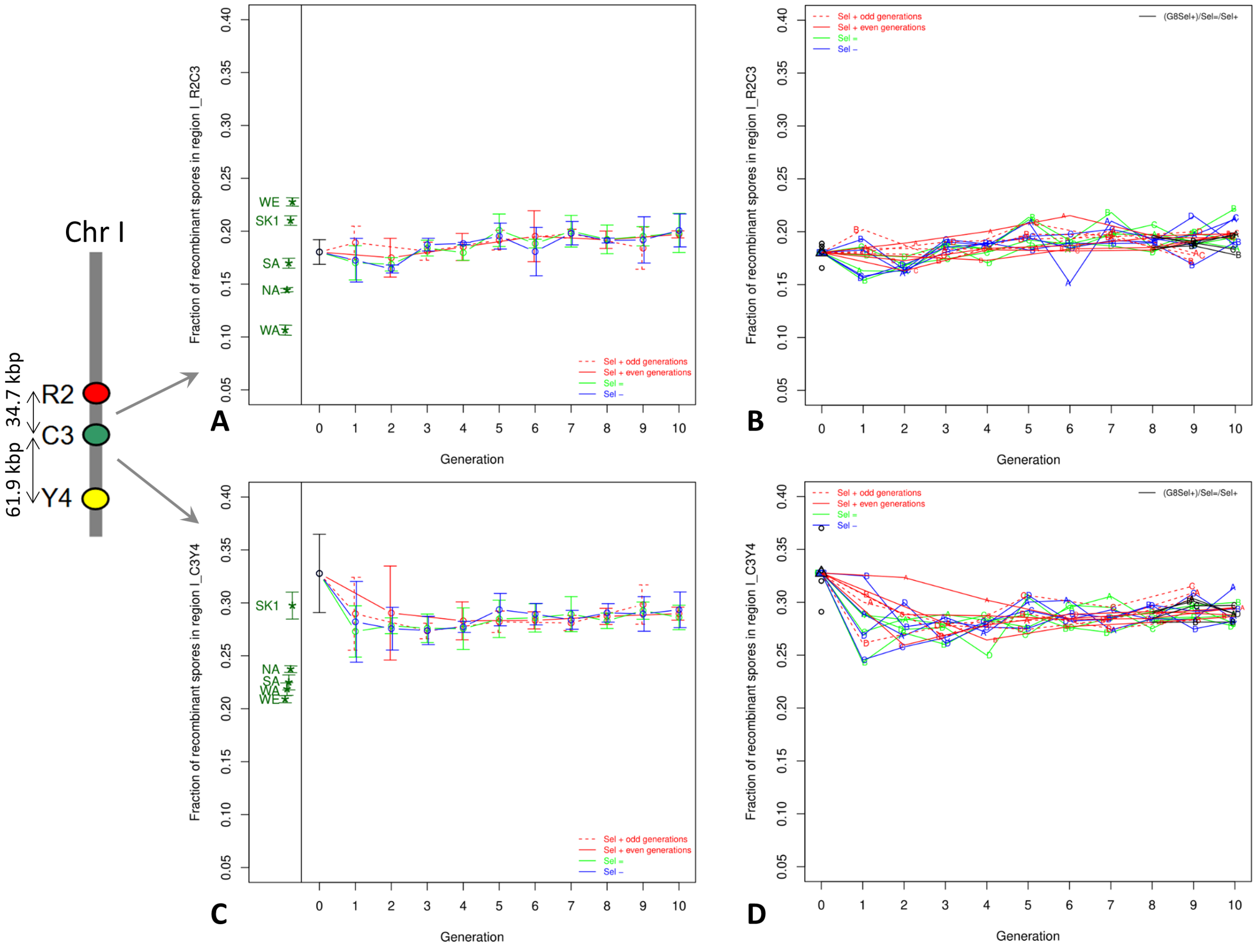

**Suppl Figure S11. Response to selection in two genomic regions on chromosome I.** Same as Figure S10 but in two regions of chromosome I: I_R2C3 **(A, B)** and I_C3Y4 **(C, D)**.
