## Supplementary material for "Response to divergent selection on meiotic recombination in Saccharomyces cerevisiae": Suppl Figure S12

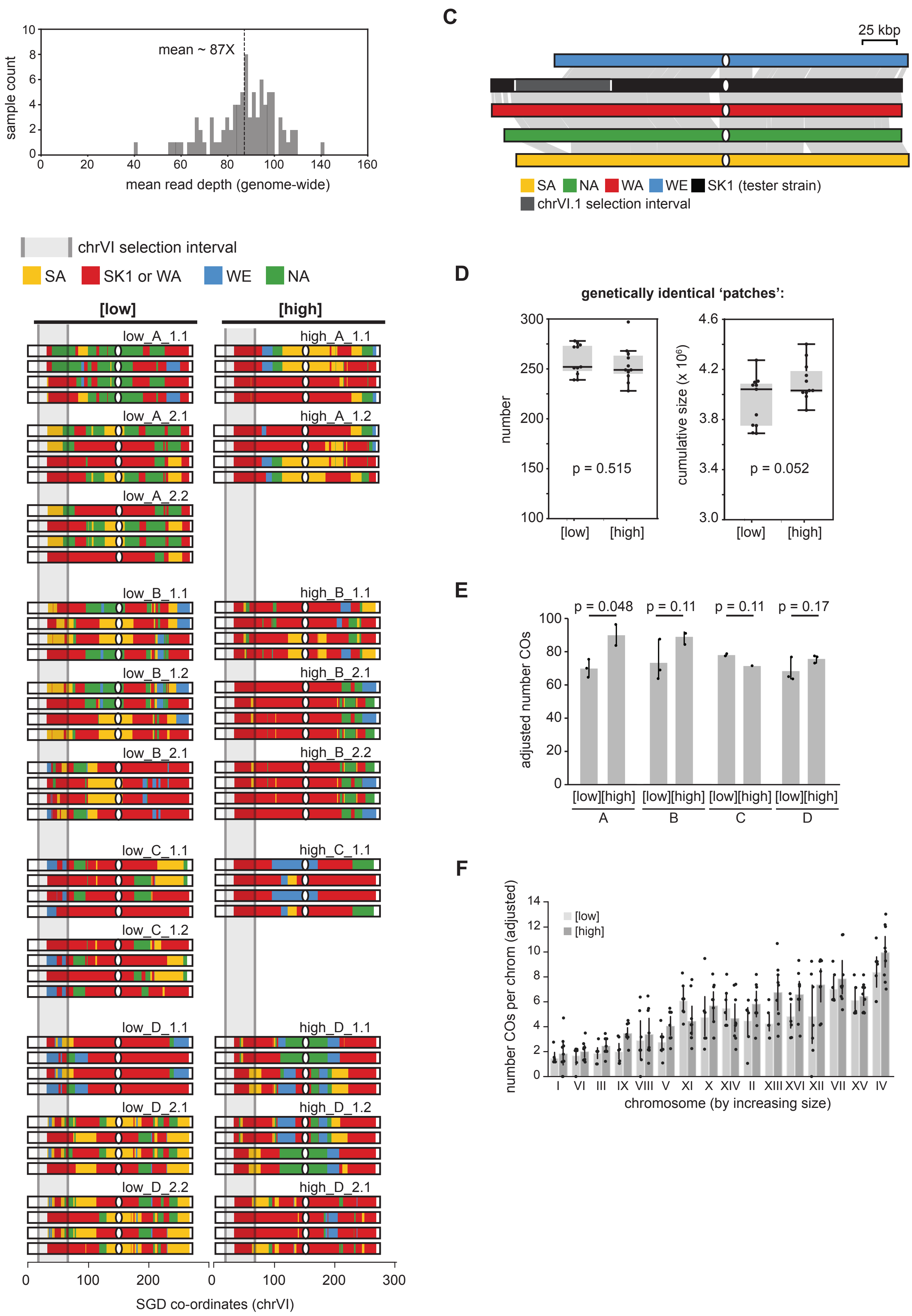

sample count
mean ~ 87X
mean read depth (genome-wide)
chrVI selection interval
SA
SK1 or WA
WE
NA
[low]
[high]
low_A_1.1
low_A_2.1
low_A_2.2
low_B_1.1
low_B_1.2
low_B_2.1
low_C_1.1
low_C_1.2
low_D_1.1
low_D_2.1
low_D_2.2
high_A_1.1
high_A_1.2
high_B_1.1
high_B_2.1
high_B_2.2
high_C_1.1
high_D_1.1
high_D_1.2
high_D_2.1
SGD co-ordinates (chrVI)
C
25 kbp
SA
NA
WA
WE
SK1 (tester strain)
chrVI.1 selection interval
D
genetically identical 'patches':
number
p = 0.515
cumulative size (x 10^6)
p = 0.052
[low]
[high]
E
adjusted number COs
p = 0.048
p = 0.11
p = 0.11
p = 0.17
[low][high]
A
B
C
D
F
number COs per chrom (adjusted)
[low]
[high]
I
VI
III
IX
VIII
V
XI
X
XIV
II
XIII
XVI
XII
VII
XV
IV
chromosome (by increasing size)

**Suppl Figure S12 Genome-wide analysis of recombination following eight generations of selection. (A):** Mean read depth for each of the 80 sequenced sergregants. **(B):** Individual genotype compositions across chromosome VI for the 9 high-recombining ([high]) tetrads and 11 low-recombining tetrads ([low]) at G8 Sel+. **(C)**: Homology plot for chromosome VI between the 4 parental genotypes (SA, NA, WA, WE) as well as the SK1 tester strain. Each grey connection bar represents a section of homology detected by mVISTA (probability threshold 0.5) with an additional filter of minimum conservation 85% and minimum block size 1 kbp. **(D):** Quantification of the number (left) and cumulative size (right) of the four-way genetically identical patches in which GCs cannot be detected, and COs cannot be precisely located. **(E):** Breakdown of the CO rate shown in Figure 4C by biological replicate. *p*-value indicates the non-adjusted result of each independent *t*-test. (F): CO quantification by chromosome, with chromosomes ordered by increasing size. *p*-value indicates the non-adjusted result of each independent *t*-test.
