## Supplementary material for "Response to divergent selection on meiotic recombination in Saccharomyces cerevisiae": Suppl Figure S13

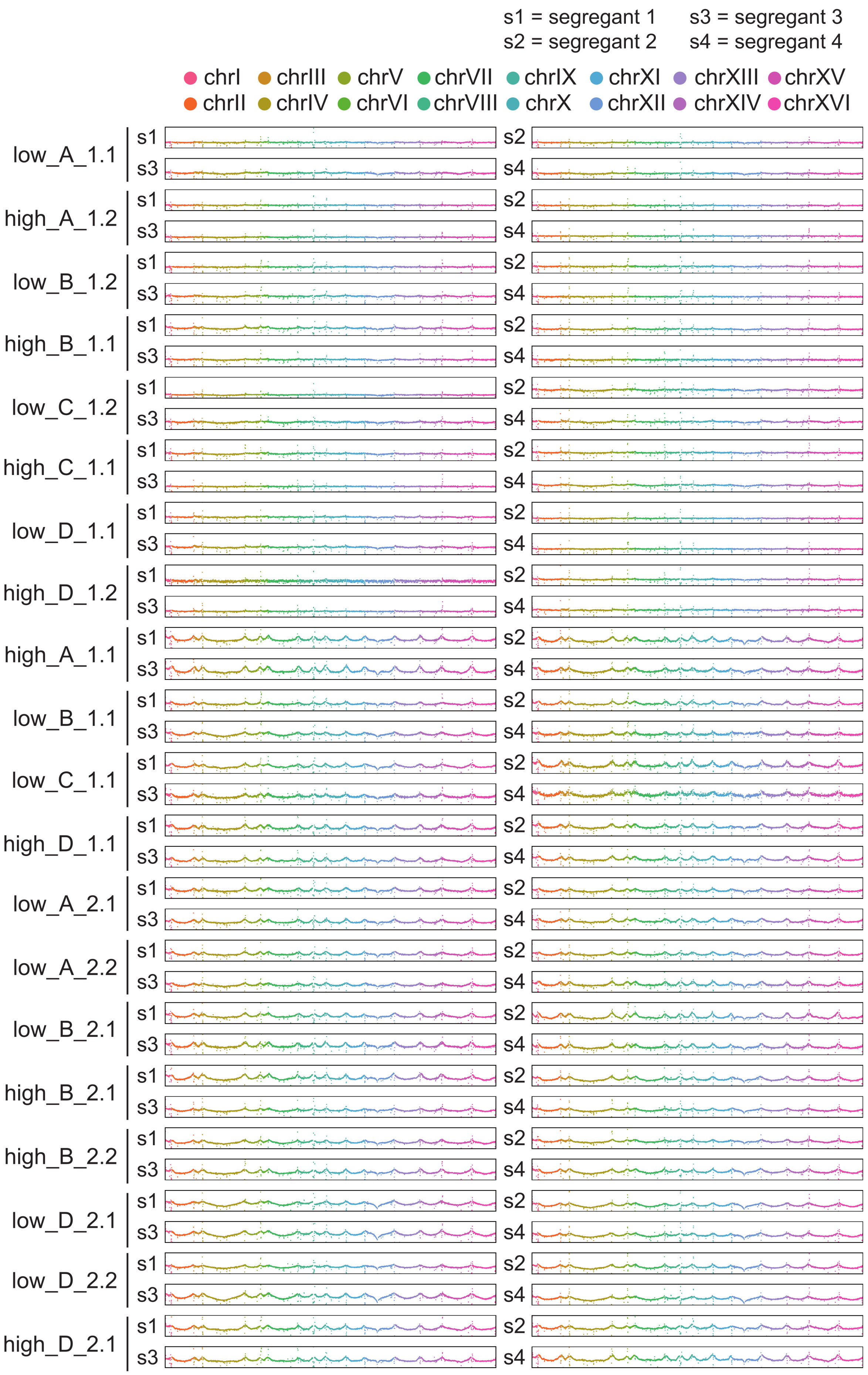

s1 = segregant 1
s2 = segregant 2
s3 = segregant 3
s4 = segregant 4
chrI chrII chrIII chrIV chrV chrVI chrVII chrVIII chrIX chrX chrXI chrXII chrXIII chrXIV chrXV chrXVI
low_A_1.1
high_A_1.2
low_B_1.2
high_B_1.1
low_C_1.2
high_C_1.1
low_D_1.1
high_D_1.2
high_A_1.1
low_B_1.1
low_C_1.1
high_D_1.1
low_A_2.1
low_A_2.2
low_B_2.1
high_B_2.1
high_B_2.2
low_D_2.1
low_D_2.2
high_D_2.1
s1
s2
s3
s4

**Suppl Figure S13 CNV analysis of all sequenced segregants**. Genome-wide read counts of 80 sequenced segregants to ensure that there are no chromosomal aneuploidies or major CNVs. Each data point represents an average read count across a non-overlapping 5-kbp window.
