## Supplementary material for "Response to divergent selection on meiotic recombination in Saccharomyces cerevisiae": Suppl Figure S14

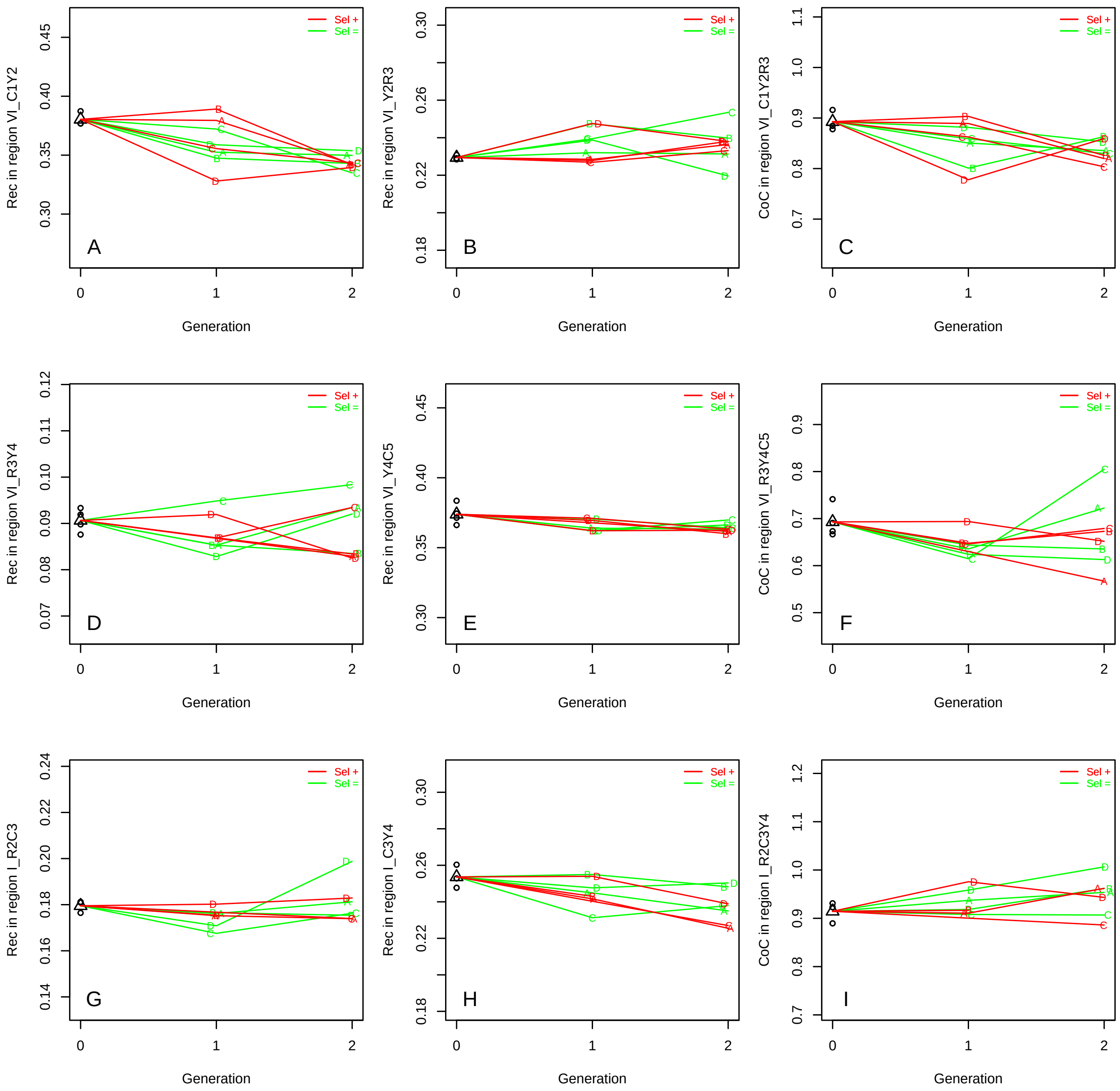


**Suppl Figure S14. Response to selection without initial genetic diversity.** Recombination rate (RR) and coefficient of coincidence (CoC) during two generations of selection for recombinant spores (Sel+, red lines) or without selection (Sel=, green lines) from a SK1 VI_C1Y2 diploid strain hemizygous for the two fluorescent markers C (yECerulean) and Y (Venus) on chromosome VI. Values of RR at each generation in intervals VI_C1Y2 where selection was applied **(A),** VI_Y2R3 **(B)**, VI_R3Y4 **(D)**, and VI_Y4C5 **(E)** of chromosome VI, and I_R2C3 **(G)** and I_C3Y4 **(H)** of chromosome I, and coefficients of coincidence (CoC) in double intervals VI_C1Y2R3 **(C)** and VI_R3Y4C5 **(F)** of chromosome VI, and I_R2C3Y4 **(I)** of chromosome I.
