## Supplementary material for "Response to divergent selection on meiotic recombination in Saccharomyces cerevisiae": Suppl Figure S15

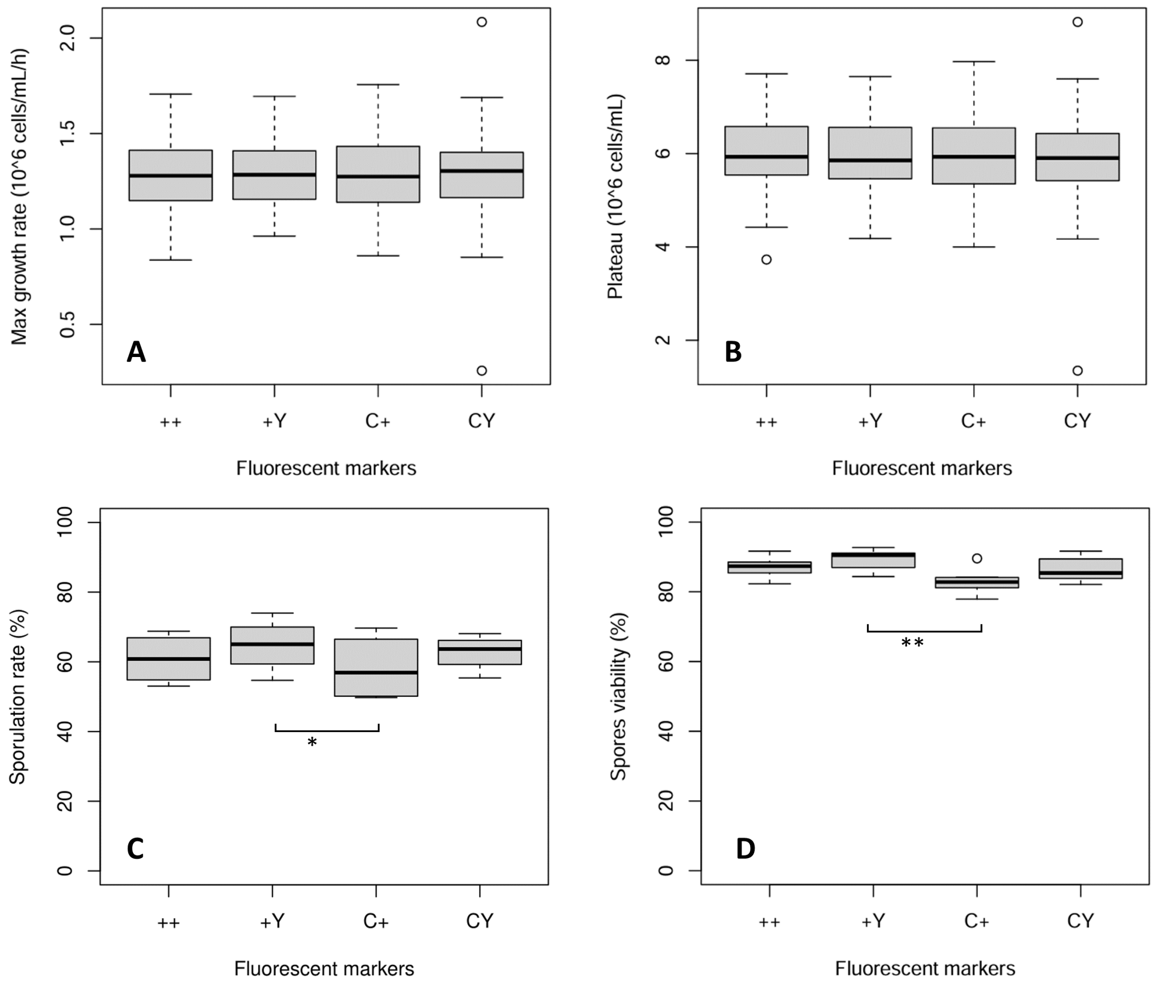


**Suppl Figure S15. Effect of fluorescent markers on growth kinetics.** Maximum growth rate **(A)**, concentration at the plateau **(B)**, sporulation rate **(C)**, and spore viability **(D)** of sub-samples (N>50,000) of population G0 sorted according to the presence of fluorescent markers. X-axis: ++, +Y, C+, and CY indicate the presence of fluorescent markers: no marker, Venus only, yECerulean only, and yECerulean and Venus, respectively. Boxplots were made from a minimum of 110 independent measurements for (**A**) and (**B**), and of 4 (**C**) or 8 (**D**) independent countings of 96 spores. Pairwise statistical comparisons shown on the graph correspond to Bonferroni-adjusted *p*-values. * <0.05, ** <0.01.
