## Supplementary material for "Response to divergent selection on meiotic recombination in Saccharomyces cerevisiae": Suppl Table S1

**Suppl Table S1**. Details of all fluorescent marker constructions inserted in the genome of the testers

Details of all fluorescent marker constructions inserted in the genome of the tester

| Chromosome | Locus # | Construct | Insertion position (SK1 reference genome from SGRP project *) | Physical distance from the previous marker (bp) | 100 bp surrounding the insertion locus |
|---|---|---|---|---|---|
| I | 2 | TDH3prom-**mCherry**-*KanMX* | 125442 | | CTTGGTAATGTTGTTAGTCTCGAGAAATGACCTTTTTTTTACCCTCAAAA**[CONSTRUCT]**AGAATGCAACACTATTAATAAACAGTACACGAAACGGATCTTTCCGTAAG |
| I | 3 | TDH3prom-**yECerulean**-*NatMX* | 160174 | 34732 | CTATATAAGGAATCGTGTTTATTAATTGAATTTATTCCGGGAATATTCAA**[CONSTRUCT]**GTTATGTATATCTCTTTTCATATTCTTAAATACACATACTCATAATATCT |
| I | 4 | TDH3prom-**YFP**-NatMX | 222040 | 61866 | ATGAAACGATTATTTTCTTGAATAATTCGTTGTCACAATTTAACAGAAAA**[CONSTRUCT]**ATTACTATCTTTTTAAGTTCACGAAGAGACTCAAACTAAGATCATGGAGA |
| VI | 1 | TDH3prom-**yECerulean**-*NatMX* | 17412 | | GCACGTGAAAACGGTGAACGTGGTTGTATACCTTCGATACGGATTTGCTT**[CONSTRUCT]**ACTAATTGAGAGCAAAATTAGTAAGCGAAATGTGAAAATTGGCTTACGAA |
| VI | 2 | TDH3prom-**YFP**-NatMX | 87335 | 69923 | CCCAGAGAGAAAAAAGGAAAAATTTAGCTATGAAACCTCAATAAGCTTTT**[CONSTRUCT]**TAATACACCAAAGATTCAAGATAAGAGCATAGAACGAACTGTAGAATAGT |
| VI | 3 | TDH3prom-**mCherry**-*KanMX* | 159705 | 72370 | TTATATTGTATTTTTTTAAAAATTCTAATGAAGTAGTTAGTGGAATGAAT**[CONSTRUCT]**GGGAAAGAGAAAACGCGTTAGGAGGGAAAAGGCTCCTTAAATAACAGTAA |
| VI | 4 | TDH3prom-**YFP**-NatMX | 198955 | 39250 | AAGGTCAAAGGGCAGATAATGGATTCCAGACTAGGAGTGCATATAAATGG**[CONSTRUCT]**ACACTTGGATTCTTCATTAGATGGTCACTGTATCAGATCTTAGCAATTTC |
| VI | 5 | TDH3prom-**yECerulean**-*NatMX* | 262866 | 63911 | ATGTGCAATTTTGCTGATGGAGCTTGCTAAAGTTGTGATATTAGTCGAGA**[CONSTRUCT]**TCCAATTAGAACGGATTAGGCGGGAATCGCTTGAGGGATCGCCAAAAATA |

Supplementary table: details of all insertion loci where constructs were inserted.

**Example**: *SK1-I-R2C3Y4* indicates a strain SK1, carrying an mCherry marker inserted on chromosome I at locus 2, yECerulean marker at locus 3, and YFP marker at locus 4.

* Liti et al, Population genomics of domestic and wild yeasts Nature 2009
http://www.moseslab.csb.utoronto.ca/sgrp/
