## Supplementary material for "Response to divergent selection on meiotic recombination in Saccharomyces cerevisiae": Suppl Table S2

### Selection of diploid strains isolated from G8 for tetrad sequencing

| Sample Name | Mean Rec AB | Direction | Nb RR Measurement | Biological Replicate | Selection Mode |
|---|---|---|---|---|---|
| VAR_PLATE_1A_VI_C1Y2_20230118_A10 | 0.165 | [Low] | 5 | RepBio A | SelPos |
| VAR_PLATE_1A_VI_C1Y2_20230118_C5 | 0.186 | [Low] | 5 | RepBio A | SelPos |
| VAR_PLATE_1A_VI_C1Y2_20230118_C9 | 0.399 | [High] | 5 | RepBio A | SelPos |
| VAR_PLATE_1A_VI_C1Y2_20230118_A5 | 0.412 | [High] | 5 | RepBio A | SelPos |
| VAR_PLATE_4A_VI_C1Y2_20230201_A11 | 0.185 | [Low] | 5 | RepBio B | SelPos |
| VAR_PLATE_4A_VI_C1Y2_20230201_E3 | 0.185 | [Low] | 5 | RepBio B | SelPos |
| VAR_PLATE_4A_VI_C1Y2_20230201_B10 | 0.374 | [High] | 5 | RepBio B | SelPos |
| VAR_PLATE_4A_VI_C1Y2_20230201_A5 | 0.403 | [High] | 5 | RepBio B | SelPos |
| VAR_PLATE_7A_VI_C1Y2_20230215_H11 | 0.160 | [Low] | 5 | RepBio C | SelPos |
| VAR_PLATE_7A_VI_C1Y2_20230215_H6 | 0.173 | [Low] | 5 | RepBio C | SelPos |
| VAR_PLATE_7A_VI_C1Y2_20230215_G11 | 0.353 | [High] | 5 | RepBio C | SelPos |
| VAR_PLATE_7A_VI_C1Y2_20230215_F4 | 0.411 | [High] | 4 | RepBio C | SelPos |
| VAR_PLATE_10B_VI_C1Y2_20230524_D2 | 0.164 | [Low] | 5 | RepBio D | SelPos |
| VAR_PLATE_10B_VI_C1Y2_20230524_F10 | 0.211 | [Low] | 5 | RepBio D | SelPos |
| VAR_PLATE_10A_VI_C1Y2_20230222_G12 | 0.378 | [High] | 5 | RepBio D | SelPos |
| VAR_PLATE_10A_VI_C1Y2_20230222_G2 | 0.381 | [High] | 4 | RepBio D | SelPos |

**Suppl Table S2.** Selection of diploid strains isolated from G8 Sel+ populations for tetrad sequencing
