## Supplementary material for "Response to divergent selection on meiotic recombination in Saccharomyces cerevisiae": Suppl Table S3

Effect of Selection mode and Biological replicate on RR measured in 770 individuals isolated from G8 populations. Pairwise Wilcoxon Test results (Bonferroni-adjusted p-values)

| Group1 | Sel− Rep A | Sel− Rep B | Sel− Rep C | Sel− Rep D | Sel+ Rep A | Sel+ Rep B | Sel+ Rep C | Sel+ Rep D | Sel= Rep A | Sel= Rep B | Sel= Rep C |
|---|---|---|---|---|---|---|---|---|---|---|---|
| Sel− Rep B | 1 | – | – | – | – | – | – | – | – | – | – |
| Sel− Rep C | 1 | 1 | – | – | – | – | – | – | – | – | – |
| Sel− Rep D | 0.0683 | 0.07 | 0.173 | – | – | – | – | – | – | – | – |
| Sel+ Rep A | <0.001 | <0.001 | <0.001 | <0.001 | – | – | – | – | – | – | – |
| Sel+ Rep B | <0.001 | <0.001 | <0.001 | <0.001 | 1 | – | – | – | – | – | – |
| Sel+ Rep C | <0.001 | <0.001 | <0.001 | <0.001 | 0.0467 | 0.328 | – | – | – | – | – |
| Sel+ Rep D | <0.001 | <0.001 | <0.001 | <0.001 | 1 | 1 | 0.0772 | – | – | – | – |
| Sel= Rep A | 0.215 | 0.515 | 0.152 | 0.00106 | 0.153 | 0.438 | 1 | 0.174 | – | – | – |
| Sel= Rep B | <0.001 | 0.00173 | <0.001 | <0.001 | 1 | 1 | 1 | 1 | 1 | – | – |
| Sel= Rep C | <0.001 | 0.00283 | <0.001 | <0.001 | 1 | 1 | 1 | 1 | 1 | 1 | – |
| Sel= Rep D | 0.00353 | 0.0253 | 0.00546 | <0.001 | 0.058 | 0.0931 | 1 | 0.0357 | 1 | 1 | 1 |

**Suppl Table S3.** Effect of selection mode and biological replicate on RR measured in 770 individuals isolated from the populations at generation G8. Bonferroni-adjusted *p*-values from pairwise Wilcoxon Test.
