## Supplementary material for "Response to divergent selection on meiotic recombination in Saccharomyces cerevisiae": Suppl Text

### Supplementary Text

ADDITIONAL METHODS

#### 1. Insertion positions of fluorescent markers used for selection

The TDH3prom-yECerulean-natMX construct (named “C1”) is inserted on chromosome VI at position 17,412 bp (S288c_reference_genome_R57-1-1). The 100 bp surrounding the insertion locus are: GCACGTGAAAACGGTGAACGTGGTTGTATACCTTCGATACGGATTTGCTT[CONSTRUCT]ACTAATTGAGAGCAAAATTAGTAAGCGAAATGTGAAAATTGGCTTACGAA. The TDH3prom-Venus-NatMX construct (named “Y2”) is inserted on chromosome VI at position 87,335 bp (S288c_reference_genome_R57-1-1). The 100 bp surrounding the insertion locus are: CCCAGAGAGAAAAAAGGAAAAATTTAGCTATGAAACCTCAATAAGCTTTT[CONSTRUCT]TAATACACCAAAGATTCAAGATAAGAGCATAGAACGAACTGTAGAATAGT.

#### 2. Sporulation of the G0 population and spores isolation

To produce isolated spores from populations, first, a vegetative liquid culture of the population was grown overnight (15 mL YPD in 50 mL tubes, 30° C, 200 rpm), then centrifuged 2 min at 2000 rpm. The supernatant was discarded, and cells were resuspended in 800 µL dd$H_2O$. Then 150 µL was spread on 4 Petri dishes with solid SPOR medium and incubated at 30° C. After ten days of sporulation, cells were picked up from the lawn by scraping one quarter of the Petri dish surface with a bent pipette tip, and resuspended in a 1.5 mL Eppendorf tube containing 750 µL dd$H_2O$ with 5 mg/mL 20T zymolyase (Euromedex, Souffelweyersheim, France) and 100 µL glass beads (0.5 mm diameter, Dutscher 67172 Brumath Ref 1606106). To disrupt tetrads, the tubes were vortexed for 30 sec three times, separated by 30 min incubations at 30° C, then centrifuged for 1 min at 13000 rpm, and the pellets were resuspended by vortexing for 2 min in 200 µL dd$H_2O$. The suspension, containing mostly vegetative cells, was discarded. Spores, which adhere to the tube plastic, were stripped by adding 600 µL dd$H_2O$ with 0.01% NONIDET NP40 (Sigma-Aldrich, Saint Quentin Fallavier, France) and vortexing 2 min, as described in Rockmill *et al.* (1991).

#### 3. Crossing the initial G0 population with the fluorescent tester

Vegetative cells of the haploid Mat-a bi-fluorescent strain SK1-VI_C1Y2 were grown overnight (15 mL YPD liquid culture in a 50 mL tube, 30° C, 200 rpm), centrifuged for 2 min at 2000 rpm, and the pellets were resuspended in 600 µL dd$H_2O$. These vegetative cells and the spores from the SGRP-4X population were crossed: both cell suspensions were mixed, vortexed for 30 sec, and centrifuged for 2 min at 2000 rpm. Pellets were resuspended in 250 µL dd$H_2O$, and 200 µL were spread on solid YPD Petri dishes and grown overnight at 30 °C. To select only diploids resulting from the desired cross, each of the four quarters of the Petri dish surface was then scraped and resuspended in 250 µL dd$H_2O$, and 200 µL was spread on SD medium without uracil and containing nourseothricin. This allowed us to select only diploids originating from crosses between a SGRP-4X spore (Ura+) and a tester cell (natMX), and none of the three other possible diploid genotypes, nor the remaining parental vegetative cells (**Suppl Figure S1**).

#### 4. Sorting of spores depending on their fluorescence status

We first selected events corresponding to the size of spores using a gate in the SSC-Height-Log *vs* FSC-Height-Log graph, then we discarded events containing more than one cell using a gate in the SSC-Height-Log *vs* SSC-Area-Log graph, see more details in Raffoux et al. (2018b). Finally, we analyzed the fluorescence intensity of each spore in yECerulean and Venus channels (excitation at 405 and 488 nm, respectively, emission at 448/59 and 526/52 nm, respectively).

Spores were then selected depending on their fluorescence status and sorted in a Petri dish containing solid YPD with chloramphenicol and glass beads (3 mm diameter). At the end of each sorting, Petri dishes were gently shaken to spread glass beads and cells on the surface of the medium.

#### 5. Sporulation and isolation of spores for G1 to G10

After 2 days of incubation at 30 °C for germination and mating of the sorted spores, a quarter of the surface of Petri dishes containing each diploid G1 population was scraped using a bent tip, resuspended in 250 µL dd$H_2O$, vortexed, and spread on Petri dishes containing SPOR medium with chloramphenicol. A second quarter of each Petri dish was also scraped, resuspended in 400 µL liquid YPD and 180 µL 50 % glycerol, vortexed, and stored at -80 °C. After 10 days of incubation at 30 °C, a quarter of each SPOR Petri dish was scraped, and we prepared isolated spores as described above, thus completing the first selection cycle. The same procedure was recurrently applied to produce further generations.

Furthermore, after the first generation, we observed that Venus fluorescence could exhibit three levels of intensity in spores (Raffoux and Falque 2024), depending on the fluorescent marker gene being present or absent in the spore genome, but also depending on the fluorescent protein being present or absent in the cytoplasm of the mother diploid cell. As an example, in Sel- experiments, some [++] spores can descend from a [++/++] diploid vegetative cell and exhibit the lowest Venus fluorescence intensity (only due to autofluorescence), but other [++] spores can also descend from a [C+/Y+] hemizygous diploid vegetative cell and thus exhibit the intermediate fluorescence intensity due to residual cytoplasmic Venus proteins. The highest fluorescence intensity class corresponds to [CY] or [+Y] spores, which do express the Venus protein. In our selection process, we discarded non-fluorescent spores descending from non-fluorescent diploid mother cells because the recombination status of such spores cannot be determined.

#### 6. Pool-phenotyping recombination in populations

As the Venus (Y) fluorescent marker can exhibit 3 fluorescence levels as mentioned above, depending on the selection mode, the positions of the gates used to select the non-fluorescent [++] spores to be crossed with the testers were not always the same: In the case of Sel- (**Figure 1B**) and Sel= (**Figure 1C**) experiments, as well as even generations of Sel+ experiments (lower part of **Figure 1A**), we sorted [++] spores belonging to the lowest intensity class of Venus fluorescence. Indeed, these spores were formed by non-fluorescent [++/++] diploids, so the fact that they may or may not have a CO between the marker loci does not affect their probability of being picked up. In the case of odd generations of Sel+ experiments (upper part of **Figure 1A**), all spores come from a diploid hemizygous or homozygous for the presence of the markers, so the Venus intensity levels showed only two peaks, and we had no other choice but to sort [++] spores descending from fluorescent diploids. As shown in the top of **Figure 1A**, half of these diploids are hemizygous for both markers [C+/Y+], so [++] spores descending from these diploids must have recombined between the markers. The other diploids, which are homozygous for the presence of one marker and homozygous for the absence of the other marker, cannot produce [++] spores.

After 2 days incubation at 30 °C for mating of the Petri dishes containing vegetative haploid tester cells and sorted spores, a quarter of each dish was scraped and resuspended in a well of a 96-wells plate containing 500 µL dd$H_2O$. We then applied to these cell suspensions two consecutive 1/8 dilutions by adding 700 µL dd$H_2O$ and transferring 100 µL to a new plate. 50 µL of these diluted cell suspensions were then deposited on Petri dishes (9 patches per Petri dish) containing solid SPOR medium with hygromycin.

After ten days incubation at 30 °C, tetrads were picked up from the lawn by scraping patches with a bent pipette tip, and suspended in a rack of 96 tubes (Macherey-Nagel Ref 740637) containing 375 µL dd$H_2O$ with 5 mg/mL 20T zymolyase (Euromedex, Souffelweyersheim, France) and 100 µL glass beads (0.5 mm diameter, Dutscher 67172 Brumath Ref 1606106). To disrupt tetrads, the plates were incubated 1 hour at 30 °C and shaken three times for 1 min at 30-minute intervals at a frequency of 23 Hz in a TissueLyser (Qiagen), then centrifuged for 5 min at 4500 rpm. Supernatants were discarded by pipetting, and the pellets were resuspended by shaking as previously in 200 µL dd$H_2O$. The suspension, mostly containing vegetative cells, was discarded. Spores, which adhere to the tube plastic, were stripped by vortexing 2 min (23 Hz) the tubes containing 400 µL dd$H_2O$ with 0.01 % NONIDET NP40 (Sigma-Aldrich, Saint Quentin Fallavier, France) following Rockmill et al. (1991). The spore suspensions were then analyzed with a CytoFLEX flow cytometer, the associated software CytExpert (Beckman-Coulter, Villepinte, France) for manual analyses, and the dedicated R package CAYSS (Raffoux and Falque 2024), which computes recombination rates in the intervals between markers, and the coefficient of coincidence (CoC) based on the two intervals delimited by the three fluorescent markers in each tester strain.

#### 7. Phenotyping recombination of isolated individuals from G8

Glycerol stocks of the G8 populations were spread on solid YPD Petri dishes with chloramphenicol and incubated for 24 h at 30 °C. Then one quarter was scraped, resuspended in YPD with Glycerol, and stored at -80 °C. Another quarter was scraped in 600 µL dd$H_2O$, and bi-fluorescent [CY] vegetative cells were sorted by FACS using a MoFlo ASTRIOS (Beckman-Coulter, Villepinte, France). During cell sorting, single bi-fluorescent cells were dropped in each well of three 96-well plates on 100 µL solid YPD. For each of the 12 populations, 288 single cells were sorted. The plates were then incubated for 48 hours at 30 °C.

Ninety-six colonies were picked for each of the four biological replicates of the Sel= populations, and 192 colonies for each of the four biological replicates of the Sel− and Sel+ populations. Each colony was then resuspended in a 96-well plate containing liquid YPD with chloramphenicol and incubated overnight without agitation at 30 °C. Plates were then centrifuged for 6 min at 4500 rpm, supernatant discarded by pipetting, and cells resuspended by adding 50 µL dd$H_2O$. Then, (1) patches of 10 µL cell suspension were deposited on the surface of a solid rectangular Petri dish containing solid SPOR medium with chloramphenicol and after 30 min drying, plates were incubated

10 days at 30 °C for sporulation, and (2) 150 µL of liquid YPD with glycerol was distributed in the remaining 40 µL of cells suspension, mixed by pipetting, and stored at -80 °C. After sporulation, cells were picked up from the lawn by scraping patches with a bent pipette tip and suspended in a rack of 96 tubes (Macherey-Nagel Ref 740637) containing 100 µL of dd$H_2O$ with 5 mg/mL 20T zymolyase (Euromedex, Souffelweyersheim, France). To disrupt tetrads, plates were incubated for 1 h at 30 °C, and shaken three times for 1 min at 23 Hz in a TissueLyser (Qiagen) separated by 30 min incubations at 30 °C, and finally centrifuged 5 min at 4500 rpm. Supernatants were discarded by flipping the plate over, and 100 µL glass beads (0.5 mm diameter, Dutscher 67172 Brumath Ref 1606106) were added, as well as 150 µL dd$H_2O$ with 0.01 % NONIDET NP40 (Sigma-Aldrich, Saint Quentin Fallavier, France). Plates were then vortexed 2 min (freq=23Hz) and stored at 4 °C before FACS analysis. Finally, the spore suspensions were analyzed, as for the pool-phenotyping, *via* a CytoFLEX flow cytometer, the associated software CytExpert, and the dedicated R package CAYSS (Raffoux and Falque 2024).

Out of 1,920 isolated diploid cells, (768 Sel+, 768 Sel− , 384 Sel=), 267 strains produced fewer than 2,000 spores (117 Sel+, 109 Sel− , 41 Sel=) and were discarded, as well as 571 which produced spores with a single fluorescence intensity peak (no segregation, due to homozygous marker) for one or both markers (217 Sel+, 175 Sel−, 179 Sel=). These observed frequencies of strains with non-segregating markers were compatible (based on exact binomial test) with the expected values (0.33 for Sel+, 0.33 for Sel− , 0.55 for Sel=; see **Suppl Figure S3**) in the case of Sel+ (observed freq=0.33, *p*-value=0.87) and Sel= strains (observed freq=0.52, *p*-value=0.30), but not for Sel- strains (observed freq=0.26, *p*-value=0.0004).

We then checked single-locus Mendelian segregation in the 1,082 remaining strains (434 Sel+ , 484 Sel−, 164 Sel=) by comparing the allele frequencies $F_C$ and $F_Y$ of C and Y markers in spores with their expected value of 0.5. Allele frequency thresholds were defined as $Thr = median(X_i) \pm 3.MAD(X_i)$ , where MAD refers to the Median Absolute Deviation, $MAD = median(|X_i - median(X)|)$. Using these thresholds, 204 strains (90 Sel+, 82 Sel−, 32 Sel=) were outliers because of non-Mendelian segregation for marker C and 206 (93 Sel+, 80 Sel−, 33 Sel=) for marker Y, so a total of 224 strains (98 Sel+, 88 Sel-, 38 Sel=) were discarded (**Suppl Figure S4**).

We also checked for two-loci Mendelian markers segregation in the 858 remaining strains (336 Sel+, 396 Sel−, 126 Sel=) by comparing the genotype frequency ratios $F_{[CY]}/(F_{[CY]}+F_{[++]})$ and

$F_{[C+]}/\left(F_{[C+]}+F_{[+Y]}\right)$ in spores with their expected value of 0.5. Using the same MAD-based genotype frequency thresholds as above, 35 strains (9 Sel+, 2 Sel−, 24 Sel=) were outliers because of non-Mendelian segregation for genotypes CY *vs* ++, and 22 strains (4 Sel+, 17 Sel−,1  Sel=) for genotypes C+ *vs* +Y, so a total of 57 strains (13 Sel+, 19 Sel-, 25 Sel=) were discarded. We observed a surplus of strains producing an excess of CY and C+ spores (**Suppl Figure S5**) as compared to other categories of non-Mendelian outliers, and that $F_{[CY]}/\left(F_{[CY]}+F_{[++]}\right)$ ratio was higher in Sel= than in the two other modes of selection (**Suppl Figure S6** ).

The numbers of strains discarded or kept at each of the different steps presented above are recapitulated in the following table.

| Sel+ | Sel- | Sel= | Category |
|---|---|---|---|
| 768 | 768 | 384 | Total isolated colonies |
| 117 | 109 | 41 | fewer than 2000 spores |
| 217 | 175 | 179 | no segregation, single fluorescence peak |
| 434 | 484 | 164 | remaining cells |
| 90 | 82 | 32 | non-Mendelian single-locus segregation C/+ |
| 93 | 80 | 33 | non-Mendelian single-locus segregation Y/+ |
| 98 | 88 | 38 | total discarded for non-Mendelian single-locus segregation |
| 336 | 396 | 126 | remaining cells |
| 9 | 2 | 24 | non-Mendelian bi-locus segregation CY/++ |
| 4 | 17 | 1 | non-Mendelian bi-locus segregation C+/+Y |
| 13 | 19 | 25 | total discarded for non-Mendelian single-locus segregation |
| 28 | 1 | 0 | outlier RR value higher than 0.5 |
| 2 | 0 | 0 | outlier RR value close to 0.5 |
| 30 | 1 | 0 | total discarded outliers for RR value |
| **293** | **376** | **101** | **ok for final RR measurement** |

In blue: remaining strains for further steps, in red: strains discarded, in grey: details of reason for discarding

#### 8. Whole-genome sequencing of an isolated diploid strain with an outlier RR value

We used Oxford Nanopore Technologies (ONT) to whole-genome sequence one diploid strain (1A-E8) isolated from the G8 Sel+ population, which had a RR of 0.491 classifying it as an outlier.

The high molecular weight DNA extraction was performed based on the protocol of Oxford Nanopore Technologies for yeast DNA, derived from a pre-existing protocol (Denis et al. 2018). The yeast strain was grown overnight in YPD at 30°C, diluted to OD600 = 0.2 in fresh YPD, and cultured to OD600 ≥ 0.7. Cells were harvested by centrifugation, washed in PBS 1X, and stored at

−80°C. Pellets were resuspended in sorbitol 1 M, treated with zymolyase (1000 U/mL) for 1 h at 30°C, and centrifuged. A lysis buffer (TrisHCl 114 mM, EDTA 115 mM, NaCl 571 mM, 1.14 % PVP40) was added, followed by SDS 10 % and RNase A, and samples were incubated for 1 h at 50°C. After the addition of TE 1X and potassium acetate 5 M, lysates were clarified by centrifugation. DNA was precipitated with isopropanol, washed with cold 70 % ethanol, air-dried, and resuspended in TE 1X overnight, then stored at −20°C.

For high-molecular-weight DNA size selection, following a protocol modified from Jones et al. (2021), the samples (3–10 µg) were diluted in TE and mixed 1:1 with 2× Size Selection Buffer (2.5 % PVP-360k, 1.2 M NaCl, 20 mM Tris-Hcl, pH 8). Samples were centrifuged at 10 000 g for 30 min, and the supernatant was discarded. The DNA pellet was washed twice with 70 % ethanol, recentrifuged at 10 000 g for 3 minutes, air-dried, and resuspended in 50 µL TE 1X at 37–50°C with gentle mixing. DNA quantity was verified using the Qubit dsDNA BR assay (Invitrogen™, Thermo Fisher Scientific).

After quantification, 1 µg of high molecular weight genomic DNA was sequenced using a Ligation Sequencing Kit (Ligation sequencing DNA V14, SQK-LSK114, 2024, Oxford Nanopore Technologies) and a MinION Flow Cell (FLO-MIN114, 2024, Oxford Nanopore Technologies). The library preparation and the sequencing were performed according to the manufacturer's instructions. The basecalling was done using Guppy software (one strand basecalling, 7.3.9+b2176f600, 2024, Oxford Nanopore Technologies).

We set out the analysis of the reads to verify that the two fluorescent cassettes were correctly positioned in the diploid genome, addressing three questions: (1) are the insertions on-target at the two intended chromosome VI loci? (2) are there any off-target insertions?, and (3) is each insertion homozygous or heterozygous? We also checked if the region between the two markers shows important sequence changes in this strain, that may explain its drastic change in RR?

ONT reads were mapped to the S. cerevisiae SK1 reference assembly (GCA_002057885.1; O'Donnell et al., 2023) with minimap2 v2.27 (Li, 2018) using the map-ont preset, then sorted and indexed with SAMtools v1.12 (Danecek et al., 2021). Reads containing the fluorescence cassettes were identified by aligning reads to the 573-bp nat1 coding sequence common to both the CFP and YFP cassettes but absent from the wild yeast genome; restricting the query to the coding sequence avoided spurious cross-mapping to the native yeast promoter and terminator. Alignments covering less than 80% of the query sequence length were discarded. Nat-1 containing reads were then mapped genome-wide, and insertion breakpoints on chromosome VI were identified from their

CIGAR strings (large internal insertions ≥2 kb, with a soft-clip fallback). These reads were then grouped based on the positions of their insertion sites. For each site, insertion size statistics (mean, range, standard deviation) were computed. To check if the insertion sites corresponded to the positions where the fluorescent cassettes were inserted, we used the 50bp flanking sequences previously chosen to insert the cassettes by homologous recombination. These sequences were searched in reads and their locations were compared to the insertion sites found. The distance between the two 50-bp flanking sequences in each read was measured, classifying the read as wild-type (~0bp gap) or knock-in (gap of approximately one cassette length). Candidate off-target alignments were evaluated by verifying that the read was flanked on both sides by genuine genomic sequence. This approach distinguishes true ectopic integration events from chimeric artifacts generated during library preparation by ligation of unrelated DNA fragments. Finally, reads spanning both integration sites were identified and inspected visually in IGV (v2.18.4).

Of 294,194 ONT reads, 95.5% mapped to the reference genome, yielding 92.7 × mean depth and 99.7% genome coverage. Forty-nine reads contained the nat1 marker sequence, of which 48 primary alignments mapped to chromosome VI and one to chromosome X. The cassette segment of this 84-kb chromosome X read was correctly bracketed by the site 1 (**Suppl Figure S9**) junction sequences of chromosome VI, identifying this read as a chimeric ligation artifact rather than a real off-target presence of the marker; no true off-target events were therefore detected.

Two integration sites were identified on chromosome VI (**Suppl Figure S9A**), both matching the 50bp flanking sequences exactly: site 1 at position 17,535 (mean insertion size=2,895 SD=20 bp, N=18) and site 2 at position 102,983 (mean size=2,814 SD=8 bp, N=16). The check based on the 50bp flanking sequences corroborated both breakpoints and revealed a knock-in:wild-type read ratio of 18:53 at site 1 *vs* 18:15 at site 2 (**Suppl Figure S9B**). Finally, a single 109kb read (d2cd89bf-2adf-4f80-8a65-8f328b0277f3; reference coordinates 14,404–123,800) was found to span both fluorescent cassettes insertion sites, carrying the wild-type allele at site 1 and the knock-in at site 2. This read was collinear with the reference, with no evidence of strand inversion or large-scale structural rearrangement between the two loci, as confirmed by visual inspection using IGV.

Altogether, these analyses showed that both fluorescent cassettes were correctly integrated at their intended chromosome VI target sites, with no detectable off-target position or rearrangement between the two sites. In agreement with the marker segregation patterns observed by flow cytometry, both insertions were confirmed to be heterozygous, with wild-type-junction reads recovered at each site. Site 1 exhibited a lower proportion of knock-in reads than expected in both independent assays, possibly reflecting locus-specific mapping or amplification bias.

#### 9. Statistical and Data analyses

Data manipulation relied on the packages dplyr (v1.2.0; Wickham et al. 2026) and stringr (v1.6.0; Wickham 2025). Figures were generated using ggplot2 (v4.0.3; Wickham 2016), with additional plotting support from ggpubr (v0.6.3; Kassambara 2026), gridExtra (v2.3; Auguie 2017), patchwork (v1.3.2; Pedersen 2025), and interactive visualizations produced with plotly (v4.12.0; Sievert 2020) and htmlwidgets (v1.6.4; Vaidyanathan et al. 2023). Document conversion used the pandoc package (v0.2.0; Dervieux 2023). Estimated marginal means and post-hoc comparisons were computed using emmeans (v2.0.2; Lenth and Piaskowski 2026). Additional statistical tests were performed using car (v3.1-5; Fox and Weisberg 2019) and lawstat (v3.6; Gastwirth et al. 2023). Data import/export used openxlsx (v4.2.8.1; Schauberger and Walker 2025) and R.utils (v2.13.0; Bengtsson 2025). The tidyverse meta-package (v2.0.0; Wickham et al. 2019) was also loaded. Sequencing alignments were visually inspected using the Integrative Genomics Viewer (IGV, v2.18.4; Robinson et al. 2011).
